# Central role and structure of the membrane pseudokinase YukC in the antibacterial *Bacillus subtilis* Type VIIb Secretion System

**DOI:** 10.1101/2020.05.09.085852

**Authors:** Matteo Tassinari, Thierry Doan, Marco Bellinzoni, Maïalene Chabalier, Mathilde Ben-Assaya, Mariano Martinez, Quentin Gaday, Pedro M. Alzari, Eric Cascales, Rémi Fronzes, Francesca Gubellini

## Abstract

Type VIIb Secretion System (T7SSb) has been recently identified in Firmicutes resembling the mycobacterial T7SSa. Despite limited sequence homology, T7SSa and T7SSb have substrates with striking structural similarities, the WXG100 proteins. Recent advances in *Staphylococcus spp*. proposed that T7SSb is involved in intra-species competition. However, the architecture and mechanism of action of this secretion complex remain largely obscure. Here, we investigate the T7SSb of *Bacillus subtilis* as a model system. We report the first evidence of *B. subtilis* ability to mediate intra- and inter-species antibacterial activity in a T7SSb-dependent manner. Then, we present the first systematic investigation of the T7SSb protein-protein network, revealing novel interactions and highlighting the central role of the pseudokinase subunit YukC in the assembly of the system. Its direct interaction with a T7SSb-secreted toxin supports its role in recruiting substrates to the secretion machinery. Finally, we solved the crystal structure of full-length transmembrane YukC defining novel structural motifs and suggesting that intrinsic flexibility modulates the orientation of the pseudokinase domains and YukC function. Overall, our results provide a better understanding on the role and molecular organisation of the T7SSb, opening new perspectives for the comprehension of this poorly characterized molecular machine.

## Introduction

The Type VII Secretion System (T7SS) was discovered in *Mycobacterium tuberculosis* from which it secretes key virulence factors required for tuberculosis infection [1]. Related genetic clusters were later identified in other Actinobacteria, in Firmicutes, and more recently in Gram-negative bacteria, opening an entire new field of investigation [2]. Based on the genetic divergence between these secretion machines in Actinobacteria and Firmicutes, they were classified in T7SSa and T7SSb, respectively. The most studied T7SSb in Firmicutes are the ones from *Staphylococcus aureus* [3], *Bacillus anthracis* [4] and *B. subtilis* [5]. T7SSa share a few homologous subunits with T7SSb, including the membrane ATPase (protein family PF01580), and the substrates belonging to the WXG100 protein family (PF06013). These substrates, about 100 residues-long, share a conserved WXG motif and form head-to-tail dimers. The high structural similarity among the WXG100 substrates suggests a similar secretion mechanism through T7SSa and T7SSb.

However, despite their relevance in human health and a large body of work, our understanding of how transport occurs through the T7SSa or T7SSb is still very limited. Only structural information on the mycobacterial T7SSa is available, including the substrate-induced oligomerization of its membrane ATPase [6], and the structures of the ESX-3 and ESX-5 T7SS membrane complexes [7–9]. Despite revealing the molecular architecture of this secretion machinery (discussed in more details below), the recent high-resolution models did not clarify the overall mechanism of secretion. Moreover, since the T7SSa and the T7SSb machines contain mostly distinct components, these systems probably evolved towards different architectures to fulfil different functions. The T7SSb system of *S. aureus* is so far the best characterized among these systems. Despite presenting unique features probably linked to its role in pathogenesis [10], the *S. aureus* T7SSb shows homologies with the *B. subtilis* one, including the genetic organization and subunit sequence similarities. The structures of soluble portions of two proteins of *S. aureus* system were obtained in the past years: the coupling protein EssC and the pseudokinase EssB. Analyses on EssC suggest that the T7SSb ATPase shares high functional and structural homology with its T7SSa counterpart. The ATPase domains are located in the cytoplasmic region of the protein. Recently, a fourth ATPase domain was found in the DUF region of the ESX-3 T7SSa EssC protein, which was proposed to be present in *S. aureus* EssC as well [11]. The N-terminus of EssC presents two forkhead-associated (FHA) domains which are unique features of the coupling proteins in T7SSb and whose role is still unknown [12, 13]. Finally, the C-terminal region presents high sequence variation in *S. aureus* of EssC [10], which would be associated to the secretion of small effectors: EsxB, EsxC and EsxD [11, 14]. However, the secretion pathway of the small effector EsxA and of the large toxin-associated complex EsaE-D-G are still unknown.

Structural information was also obtained for the pseudokinase EssB, the homologue of *B. subtilis* YukC, from both *S. aureus* and *Geobacillus thermodenitrificans* [15, 16]. Based on this model, a possible role of the T7SSb pseudokinase in signal transduction and/or in mediating the interaction with other proteins of the systems was suggested. Notwithstanding the presence of FHA domains known to interact with kinases and pseudokinases [17], such interaction was not yet demonstrated in T7SSb machineries. The *S. aureus* EssB pseudokinase was also proposed to interact with the multispanning membrane subunit EsaA [18] as well as with other T7SSb components [19]. Despite these results, the full-length structure of a T7SSb pseudokinase is still missing and its role remains unknown.

From a functional point of view, a highly diversified range of functions was shown for T7SSb in Gram-positive bacteria, such as DNA conjugation, maintenance of membrane integrity, phage resistance, virulence or bacterial competition [20–24]. In particular, in the Firmicutes pathogenic species *S. aureus* and *B. anthracis*, T7SSbs were known to play a role in persistence [3, 4, 25]. However, it was only recently that the T7SSb function in intraspecies competition was revealed in *S. aureus* and *S. intermedius* [21, 26]. Nonetheless, major differences in T7SSb-mediated bacterial competition were reported in these studies including the nature of the secreted toxin (*i.e*., belonging or not to the ‘LXG’ family). In addition, the mechanism of toxin translocation, which remains unknown, seems to have different molecular determinants. While the *S. aureus* EsaD toxin contacts the ATPase through a specific ‘adaptor protein’ EsaE [21], different WXG100 proteins were proposed to target cognate LXG toxins to the T7SSb in *S. intermedius* [26]. Altogether, those differences in the toxins identified, in their binding partners as well as in their mechanism of competition (contact-dependent or - independent) raised the hypothesis that the T7SSb could secrete different toxins using different mechanisms.

Despite these advances in understanding the potential roles of T7SSbs, the available information on subunits interactions, structures and mechanism of substrate transport is still very limited. Moreover, in non-pathogenic bacteria such as *B. subtilis*, the role of T7SSb was still unknown. Interestingly, no EsaD homolog was identified in *B. subtilis*. Instead, six genes encoding proteins of the LXG toxins family have been identified in *B. subtilis* [27], and five of these proteins were toxic when overproduced in *E. coli* [28].

Here we present an extensive structure-function analysis of T7SSb in *B. subtilis* to understand its global organisation and mechanism of substrate secretion. We illustrate the T7SSb-dependent role in intra- and inter-species bacterial competition in *B. subtilis*. The protein-protein interaction network was investigated through bacterial double-hybrid and copurification assays revealing the central role of the YukC pseudokinase in the assembly of this secretion system. Finally, we present the crystallographic structure of the full-length transmembrane protein YukC, which highlights a novel structural arrangement corroborated by *in vivo* analysis. Altogether, this work opens new perspectives on the understanding of the role and the molecular organization of T7SSb.

## Results

### T7SSb-dependent bacterial competition in B. subtilis

Recent evidence that the *S. aureus* and *S. intermedius* T7SSbs are involved in intra-species bacterial competition [21, 26, 27] opened the possibility that they could mediate similar functions in non-pathogenic Firmicutes. To investigate this possibility, we tested the ability of the *B. subtilis* T7SSb to mediate intra-species killing in a contact-dependent manner. To this purpose, we engineered a *B. subtilis* 168 ‘prey’ strain lacking one of the LXG toxin-antitoxin pairs, YxiD-YxxD [28, 29]. YxiD has cytotoxic activity when overproduced in *E. coli* and is neutralized when bound to its cognate YxxD antitoxin [28]. We reckoned that the absence of the antitoxin YxxD would leave the *B. subtilis ΔyxiD-yxxD* (*BsΔyxiD-yxxD*) strain without defense against the specific LXG toxin YxiD, causing growth inhibition or cell death in the presence of wild-type *B. subtilis* 168 able to secrete YxiD. The *yxiD-yxxD* cassette was replaced by the GFP-encoding gene, in order to monitor bacterial competition through fluorescence. After overnight incubation in contact with wild-type *B. subtilis*, the *Bs*Δ*yxiD-yxxD* fluorescence was very low (Figure 1A upper panel, left spot), demonstrating that this strain has been outcompeted by the wild-type strain. By contrast, deletion of T7SSb genes in the attacker resulted in survival of the prey, as observed with the fluorescence of the spots (Figure 1A, upper panel). The absence of outcompetition was not due to a growth defect of the T7SSb mutants as these cells showed growth kinetics comparable to the parental wildtype strain in nutrient-rich or nutrient-limiting media [30]. We concluded that the cytotoxic effect of the attacker on the prey depends on a functional T7SSb. The competition was further quantified by numbering the Colony Forming Units (CFUs). Indeed, the number of Bs*ΔyxiD-yxxD* prey cells surviving after an overnight contact with the wild-type *B. subtilis* was two orders of magnitude lower than when in contact with the strains carrying an impaired T7SSb (Figure 1A, lower panel). Quantification of the GFP signal (normalized to the OD600) confirmed the visual result (Figure 1 - figure supplement 1A). To exclude possible polar effect of the deletions, we checked the production of YueB, which is encoded by the fifth gene of the *yuk/yue* operon, and serves as receptor for phage SPP1 [31]. We observed that the *B. subtilis* 168 strains Δ*yukE*, Δ*yukD*, Δ*yukC* and Δ*yukB* were lysed by SPP1, demonstrating that these deletions have no effect on YueB production and likely on the others T7SSb subunits (Figure 1-figure supplement 1B).

**Figure 1.**
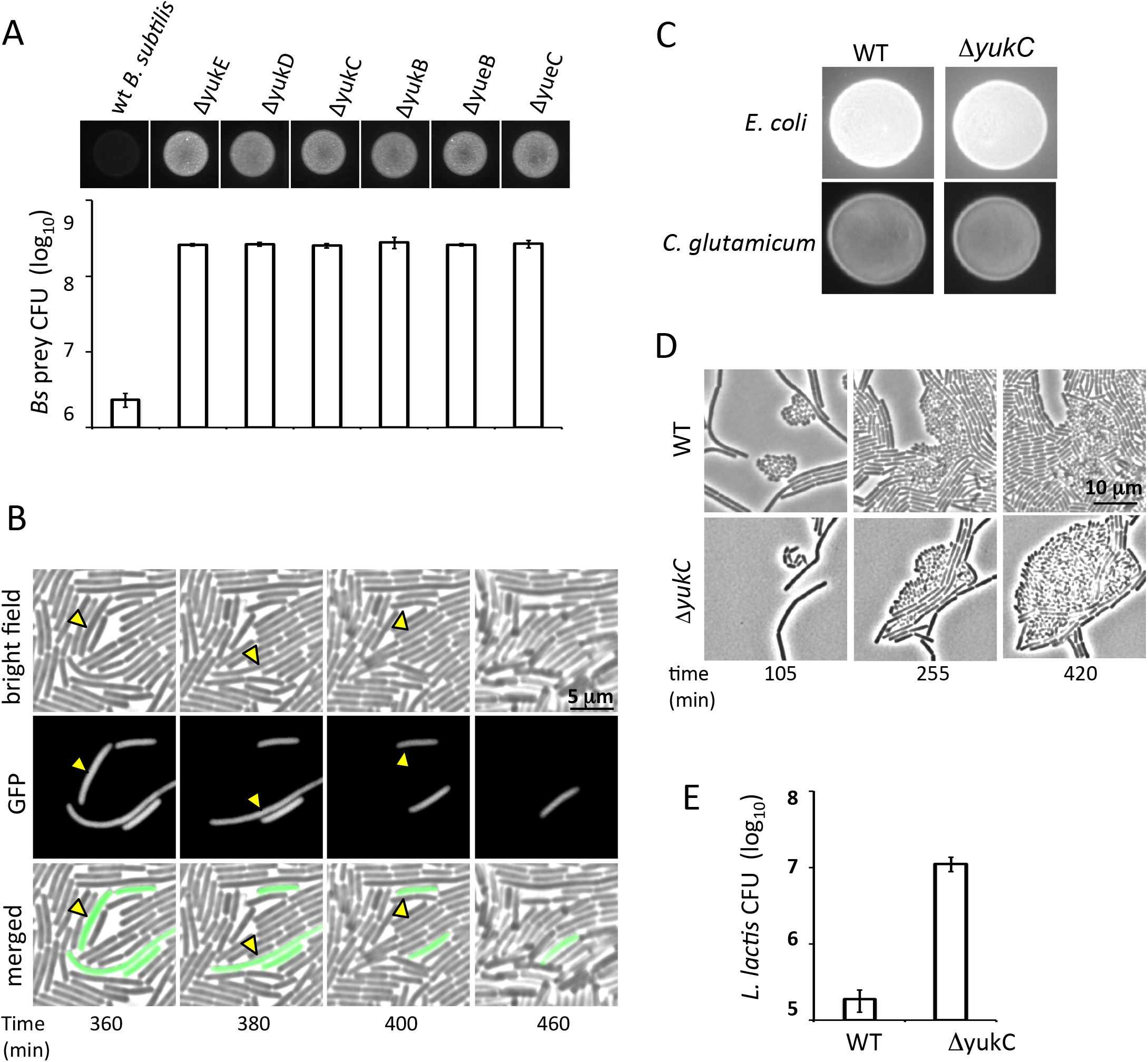
*B. subtilis* T7SSb-dependent competition assay. **A)** Competition assays using the fluorescent strain *B. subtilis ΔyxiD-yxxi::gfp* as a prey. As attackers we used the *B. subtilis 168 CA* ‘wild type’ (WT) strain and its derivative six strains deleted in a T7SSb gene each. Upward: fluorescence of bacterial spots on LB agar plate after overnight competition. In the presence of the attacker strain carrying a wild-type T7SSb operon the GFP fluorescence of the ‘prey’ disappeared (left). Instead, when strains carrying single-gene deletions in the *yuk* operon were employed, GFP fluorescence was still observed in the prey after overnight incubation. Bottom: Survival of prey cells was evaluated by the ability to form colonies on a selective media containing Spectinomycin. The CFU values are reported for each assay using a different ‘attacker’ strain as for the correspondent experiments shown above. **B)** Frames from Video 1 showing disappearance of prey cells from the bright field, and of their fluorescence from GFP and from the composite images. Samples of disappearing cells are highlighted with yellow arrows. **C)** Fluorescent spots showing the results of the competition assay against didermal bacteria *E. coli* and *C. glutamicum*. Similar fluorescence intensity is observed after overnight competition using *B. subtilis* WT and its mutant ΔyukC, suggesting no T7SSb-dependent killing of *B. subtilis* on these bacteria. **D and E)** Bacterial inter-species competition assay using *L. lactis* as ‘prey’. **D)** Frames from the live-imaging of the contactdependent competition assay showing the T7SSb-dependent killing of the monoderm Gram positive bacteria *L. lactis* by *B. subtilis* (Videos 3 and 4). **E)** Surviving *L. lactis* cells were selected on a solid media supplemented with chloramphenicol. The different survival rate in the presence of the WT and the Δ*yukC B. subtilis* strains suggest a T7SSb dependent-competition.

Altogether, these data confirmed the T7SSb-dependent ability of *B. subtilis* to kill cells of the same species in a contact-dependent manner.

To better document this mechanism of competition, we recorded the live interaction between *B. subtilis* prey and attacker cells by fluorescence microscopy (Figure 1B, Video 1). Figure 1B shows that isolated fluorescently labeled prey cells surrounded by wild-type attackers lysed. At the end of the Video 1, corresponding to the overnight incubation (about 14 hours), the overall fluorescence is confined to small patches of prey bacteria. By contrast, the fluorescent prey grew steadily without any noticeable killing event when incubated with BsΔ*yukC* attacker cells (Figure 1B and Video 2).

To determine whether the *B. subtilis* T7SSb also mediates inter-species competition, we tested the ability of *B. subtilis* 168 to compete against Gram-positive bacteria (*Lactococcus lactis subsp. cremoris* NZ9000), actinobacteria (*Corynebacterium glutamicum*) and proteobacteria (*Escherichia coli* K12 [32]). While we did not observe antibacterial activity against fluorescently labeled *C. glutamicum* and *E. coli* (Figure 1C), the survival of *L. lactis* cells was significantly affected by *B. subtilis* in a T7SSb-dependent manner (Figures 1D, 1E and Video 3). As shown in Fig. 1D, round-shaped *L. lactis* cells stopped growing after 4 hours of incubation with wild-type *B. subtilis* cells (Video 3), but continued to grow up to 9 hours when in contact with *Bs*Δ*yukC* attackers (Figure 1D and Video 4). Interestingly, we observed that *Bs*Δ*yukC* cell growth is inhibited by *L. lactis*. This inhibition may be due to the production of bacteriocins by *L. lactis* [33, 34]. Therefore, a functional T7SSb is not only required to attack other bacteria, but also serves as a counterattack mechanism against armed competitors.

### Interaction network suggests a central role for YukC in B. subtilis T7SSb

Even though limited data on subunit interactions are available on the *S. aureus* T7SSb [18, 35, 36], a full picture on the organization of this secretion machine in Firmicutes is still missing. In order to shed light onto T7SSb organization, we tested binary interactions between all its subunits.

First, we screened T7SSb subunits interactions by bacterial two-hybrid (BACTH) assay [37]. Target proteins were fused to both the adenylate cyclase T18 or T25 fragments either at their N- or C-terminus to circumvent possible steric hindrance on a specific side of the proteins (Table S2). The results of the BACTH assays are reported in Figure 2A and summarized in Figure 2B. All *B. subtilis* T7SSb proteins interacted with at least another subunits and/or with themselves, indicating that all of them were produced and properly folded in the *E. coli* strain used for the BACTH assay. We detected self-interactions for the YukC, YukB and YueB subunits, as well as for the YukE WXG100 substrate, in agreement with published data on homologous proteins [12, 15, 35, 36, 38]. As expected from the predicted topology of YukC, only the constructs having the BACTH tags (T18 and T25) at the N-terminus interacted (Figure 2 - figure supplement 1B). By contrast, no oligomerization was observed for YueC and YukD (Figure 2A). Pair-wise tests detected several interactions, including YukD-YueC and YukB-YueB. Noticeably, YukC appears to interact with all the other proteins encoded on the *yuk* operon, with the YukC-YukD interaction giving a weak but reproducible positive result (Figure 2-figure supplement 1C). This suggests a central role for the subunit YukC in the assembly of the T7SSb complex. The fact that YukE, YueB and YukB interact with YukC independently of the position of their T18 or T25 domains suggests that these proteins do not need free N- or C-terminus to contact YukC (Figure 2-figure supplement 1D).

**Figure 2.**
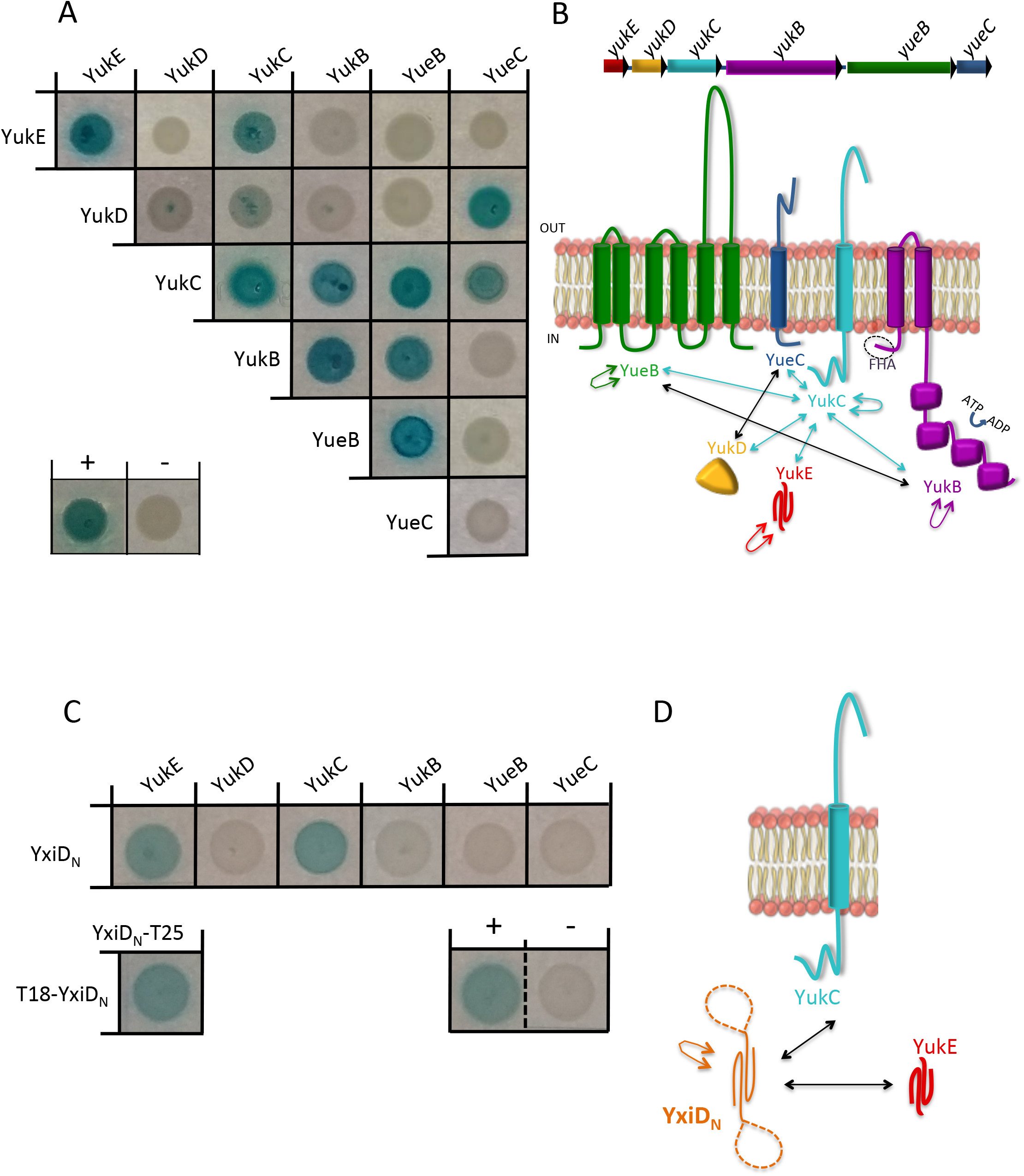
Binary protein interactions within *B. subtilis* T7SSb. **A)** Blue spots of *E. coli* cells shows interaction between the BACTH-tagged *Bs*T7SSs subunits as indicated in the scheme. Representative positive and negative controls are shown labeled respectively as (+) and (-). **B)** Top: *B. subtilis* T7SSb operon gene organization. Bottom: schematic of the binary interactions in T7SSb. The arrows are color-coded as follows: in cyan YukC interactions, in black interactions between others T77Sb subunits. Self-interactions are colored as each specific subunit. **C)** BACTH assay showing direct interaction of YxiD N-terminal region (YxiD_N_) exclusively with YukE and YukC among all the T7SSb subunits. YxiD_N_ also exhibited self-interaction. The relative controls are reported below. **D)** Schematic of YxiD_N_ interactions.

Deepening the analysis of YukC-YukB assembly, we showed that YukC interacts with the N-terminal region (residues 1-256) of the YukB ATPase (Figure 2-figure supplement 1E). This region, as well as the homologous coupling protein of others T7SSb includes two FHA domains, which are not found in T7SSa coupling proteins (Figure 2-figure supplement 2).

We then used the BACTH assay to investigate interactions between T7SSb subunits and one of the LXG toxins. As a model, we picked the previously characterized YxiD toxin, which ribonuclease activity appears localized in the YxiD C-terminal domain [28]. Therefore we used the construct YxiD_N_ corresponding to the first 97 amino acids, to avoid toxicity. This region is proposed to adopt a coiled coil fold similar to the one observed in WXG proteins. YxiD_N_ interacted with itself, as well as with the YukC T7SSb subunit and the YukE WXG100 protein (Figure 2C and 2D). This result represents the first evidence of where LXG toxin would contact the T7SSb complex (YukC) and is in agreement with previous data supporting the existence of LXG-WXG100 complexes [26].

Pair-wise interactions between YukC and the T7SSb subunits YueB, YukE and the YukB ATPase (*via* its FHA domains), were then assayed by co-purification using affinity chromatography (Figure 3). We confirmed interactions between YukC and YueB (Figure 3A), and YukC and the YukB N-terminal domain (YukBΔ_256_; Figure 3C). In these cases we detected these subunits using immunoblotting against affinity tags. Since the His_6_ tag got cut out from YukE during its co-purification with YukC, we identified their bands by SDS-PAGE (Figure 3B) coupled with mass spectrometry (MS) analysis. In addition, tackling the details of YukC-YukB interaction, we obtained a stable complex between the YukB FHA domain (YukBΔ_256_) and the isolated YukC pseudokinase domain (YukCΔ_217_ including residues 1-217), supporting the conclusion that the YukC-YukB interaction occurs through these regions (Figure 3D).

**Figure 3.**
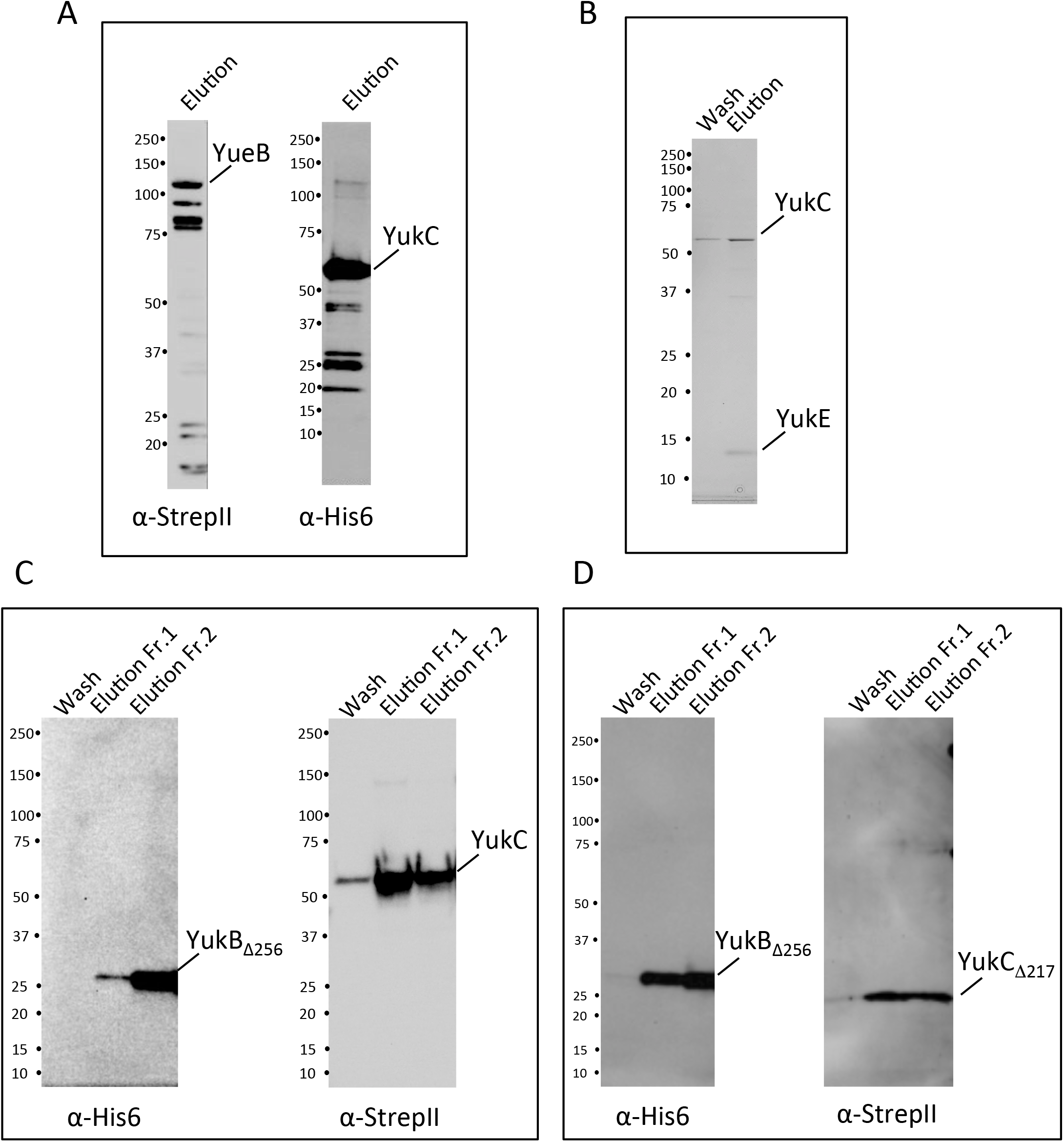
Co-purification of YukC with its interactions partners. **A)** Immunodetection of the co-purified *YueB.strep* (α-StrepII, left) and *YukC.his* (α-StrepII, *right*). **B)** SDS-PAGE migration of the eluted sample from the co-purified protein complex composed by *YukC.strep* and *YukE.his*. **C)** Immunoblot of the eluted proteins from *YukB_Δ256_.his* (FHA domains) – *YukC.strep* co-purification. The proteins have been detected using primary antibodies α-His6 (left) or α-StrepII. **D)** Immunodetection of the co-purified *YukC_Δ217_.strep* (pseudokinase domain) (α-StrepII, right) and *YukB_Δ256_.his* (α-His6, left).

### The YukC structure

The presence of multiple interactions with the others T7SSb components highlighted the central role of YukC. Previous studies yielded models of the soluble domains of YukC homologs in *G. thermodenitrificans* and *S. aureus* [15, 16]. However, the lack of the central transmembrane region in these constructs left uncertainties on the structural and oligomeric organization of the T7SSb pseudokinases, limiting structure/function interpretation.

To investigate its structure, the full-length YukC from *B. subtilis* was purified from *E. coli* generating slow-growing crystals showing poor diffraction (20 Å). This was significantly improved using the YukC_413_ construct lacking the last 38 amino acids of YukC, which are highly positively charged. The SDS-PAGE migration of purified YukC_413_ differed from the full-length YukC probably due to the stretch of positive residues lacking at its C-terminus (Figure 4-figure supplement 1A).

We solved the structure of YukC_413_ by molecular replacement on a 2.6 Å resolution dataset (PDB ID 6Z0F, Figure 4, Table 1), using the coordinates of the soluble domains from *G. thermodenitrificans* EssB as the search models (PDBs 2YNQ and 4ANO). In the crystal, YukC_413_ organizes as a pseudo-symmetrical dimer with chains A and B revolving around a virtual central axis. The last 27 residues of the YukC^413^ construct could not be traced neither in chain A (384 residues) nor in chain B (377 residues), due to the lack of supporting electron density. In the crystal packing, two YukC_413_ dimers associate in an antiparallel arrangement, with the N-terminal region of one dimer contacting the C-terminus of the other (Figure 4-figure supplement 1B) in a non-physiological organization.

**Figure 4.**
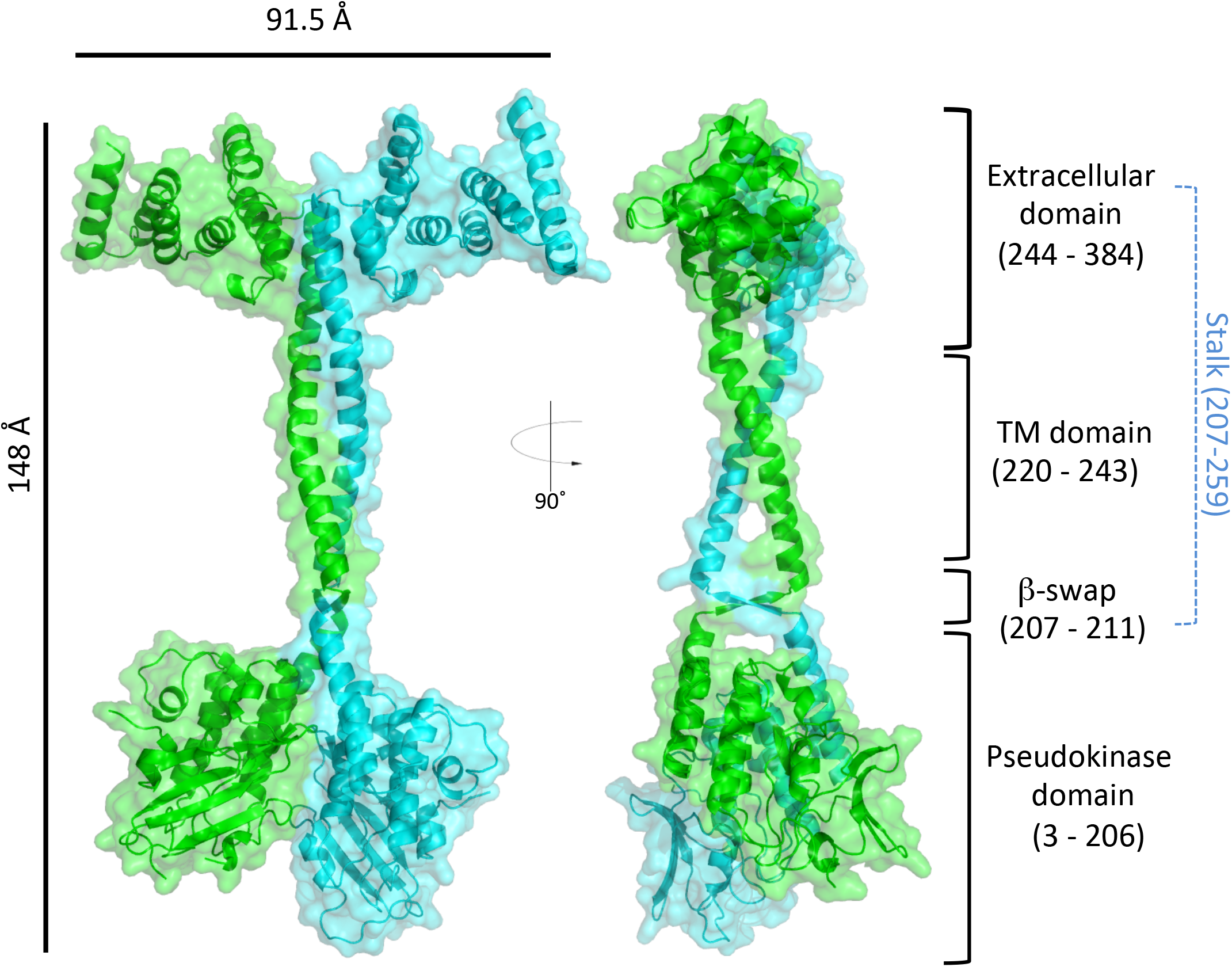
Structure of YukC_413_. Atomic model of YukC_413_ crystallographic structure (PDB ID 6Z0F). Left: front view. Right: side view. YukC_413_ dimer is rendered in cyan (chain A: 384 residues) and green (chain B: 377 residues). Four major domains of the proteins are indicated on the right: the pseudokinase domain (residues 3-206), the *β*-swap domain (residues 207-211) the transmembrane (TM) region (residues 220-243) and the extracellular domain (residues 244-384). The central stalk is indicated in blue, going from the intracellular to the extracellular region of YukC (residues 206-259).

**Table 1.**
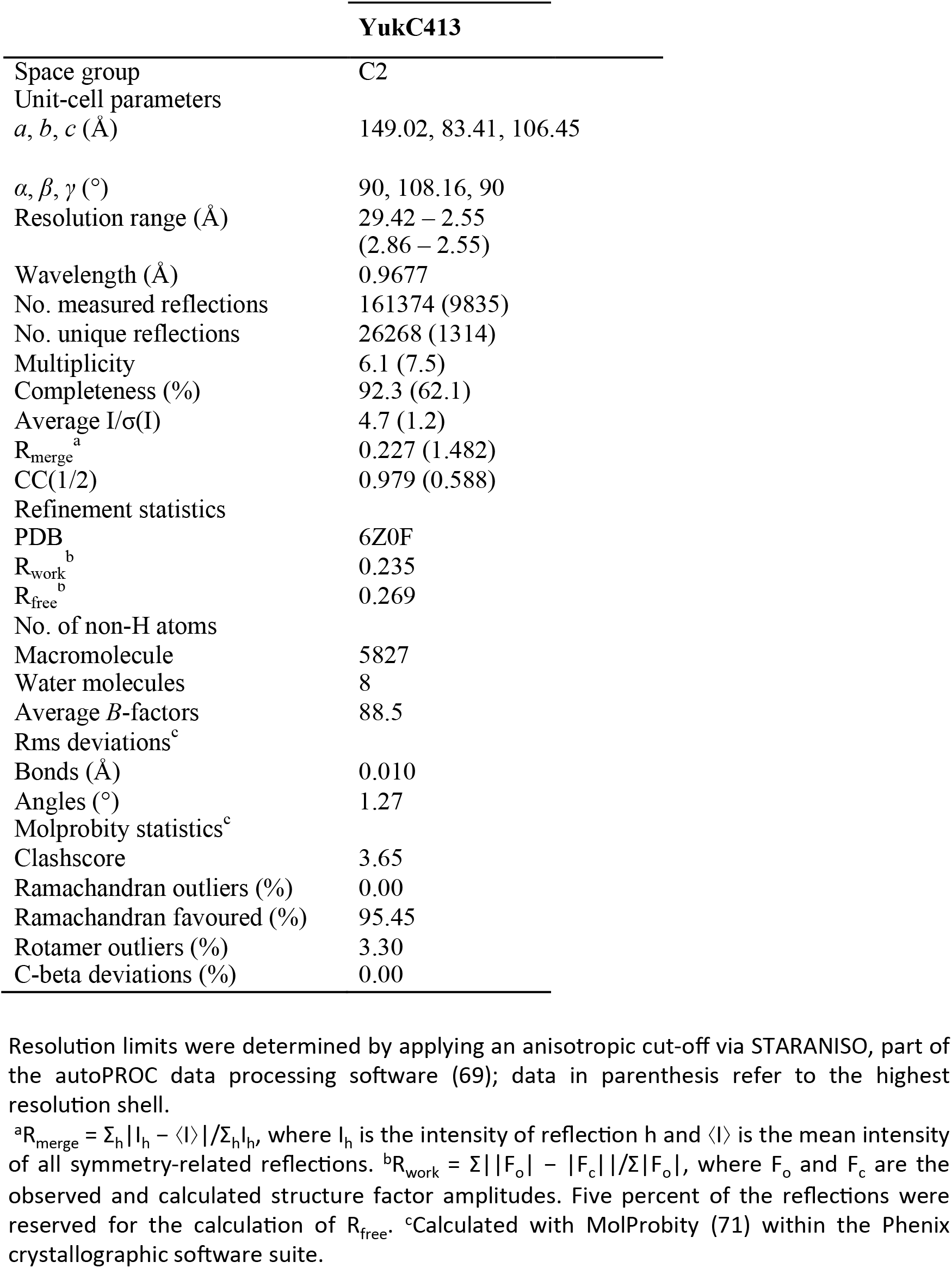

The YukC_413_ structure can be divided in four main regions, going from the N- to the C-terminus: the pseudokinase domain (residues 3-206), the ‘β-swap’ domain (residues 207-211), the transmembrane (TM) domain (residues 220-243), and the extracellular domain (residues 244-384) (Figure 4). The TM region was defined based on the hydrophobicity profile (Figure 4-figure supplement 1C), in accordance with the TMHMM prediction algorithm. A central ‘stalk’ crosses the different domains, starting at the β-swap and ending on the extracellular portion of YukC (residues 207-259).

### The extracellular domain

The extracellular domains (residues 244-384) present an all-α-helical organization, the structure of which is not reminiscent of other proteins or domains of known function (DALI analysis at http://ekhidna2.biocenter.helsinki.fi/dali/). Overall, the extracellular domain of YukC appears highly similar to that of *G. thermodenitrificans* EssB (*Gt*EssB, rmsd = 1.5) [15] (Figure 4-figure supplement 2A). It is composed mainly by 8 α-helices interacting in coiled-coil motifs and extending approximately 40 Å from the membrane plane. In the extracellular region, the YukC_413_ monomers interact at the level of the α-helix 1 and the loop between the α-helices 4 and 5 (Figure 4-figure supplement 2B) contributing to YukC dimer stabilization. We tested if this region might be involved in the interaction with YueB, which is the only T7SSb subunit having a large extracellular domain. Our bacterial two-hybrid results showed that YukC still interact with a variant of YueB missing the 90% of its extracellular domain (YueBΔ31-817, Figure 4-figure supplement 2C). Since the other loops of YueB are quite short, this result is in agreement with the proposal that YukC-YueB homologs interact through their TM regions in *S. aureus* [18].

### The pseudokinase domain

Structural comparison between the pseudokinase domains of YukC and the homologous EssB from *S. aureus* (PDB: 4ANN) and *G. thermodenitrificans* (PDB: 4ANO) indicate a closer similarity with the last one (Figure 4 - figure supplement 3A). This reflects the sequence identity percentage with YukC (15 % for *S. aureus* and 43 % for *G. thermodenitrificans*). Nevertheless, the pseudokinase domain (described in detail in [15]) exhibit an overall high similarity in the three models.

Additionally, the YukC_413_ structure allowed identifying the interface between the pseudokinase domains in the dimer, while in previous structures they were crystallized as monomers. Their interactions are mainly mediated by the α-helices α4 and α8 (K^83^-Q^99^ and Y^186^-T^206^ respectively), the region including β strands β5 and β6 (I^114^-F^124^) and the ‘connecting loop’ (CL) going from P^68^ to A^74^ (Figure 4 - figure supplement 3B). Despite a number of intermolecular salt bridges and hydrogen bonds formed between the two pseudokinase domains, they do not appear contributing to the stability of the dimer (see below).

In order to better characterize the pseudokinase region of YukC, we performed a sequence alignment comparing *B. subtilis* YukC and its *S. aureus* and *G. thermodenitrificans* EssB homologues, with known kinases or pseudokinases identified as best matches on a DALI search [39]. These include the eukaryotic kinases Sirk1, TNNI3K, Pim1 and Bsk8, the bacterial Ser/Thr kinase PknB [40], and the MviN pseudokinase [41] (Figure 4-figure supplement 4A). The most important motifs characterizing Hanks-type kinases, such as the glycine-rich P-loop coordinating the ATP β- and γ-phosphate, the “VAIK” motif (which conserved lysine moves toward the glutamate during the hydrolysis), the catalytic loop, the Mg-binding loop and the activation loop [40], are partially missing in the pseudokinases. The structural comparison of YukC_413_ with PknB in complex with ATP (PDB 1O6Y) (rmsd 4.2 Å for their pseudokinase domains) shows that YukC has an additional α-helix in the ATP-binding pocket, with a large side-chain (F^26^) partially occluding the ATP binding site (Figure 4-figure supplement 4B). In agreement with this observation, microscale thermophoresis assays showed that YukC is unable to bind ATP *in vitro*, in contrast to PknB (Figure 4-figure supplement 4C). Taken together, the sequence and structural analysis as well as the *in vitro* ATP-binding assays suggest that YukC, and likely the other T7SSb-associated pseudokinases, are unable to bind (and *a fortiori* to hydrolyze) ATP.

### The stalk and the ‘β-swap’ domain

The analysis of the YukC dimer interface using the PISA web tool (http://www.ebi.ac.uk/pdbe/pisa/) [42] indicated that the major contribution to surface dimerization comes from YukC middle region including the β-swap and the TM domain (Figure 5A). While the values of interacting surfaces are similar for the extracellular and the pseudokinase domains, their contribution to the dimer stability is significantly different, as indicated by their Δ^i^Gs values. Overall, these values show that YukC dimerization is supported by the intermolecular interactions occurring along the stalk.

**Figure 5.**
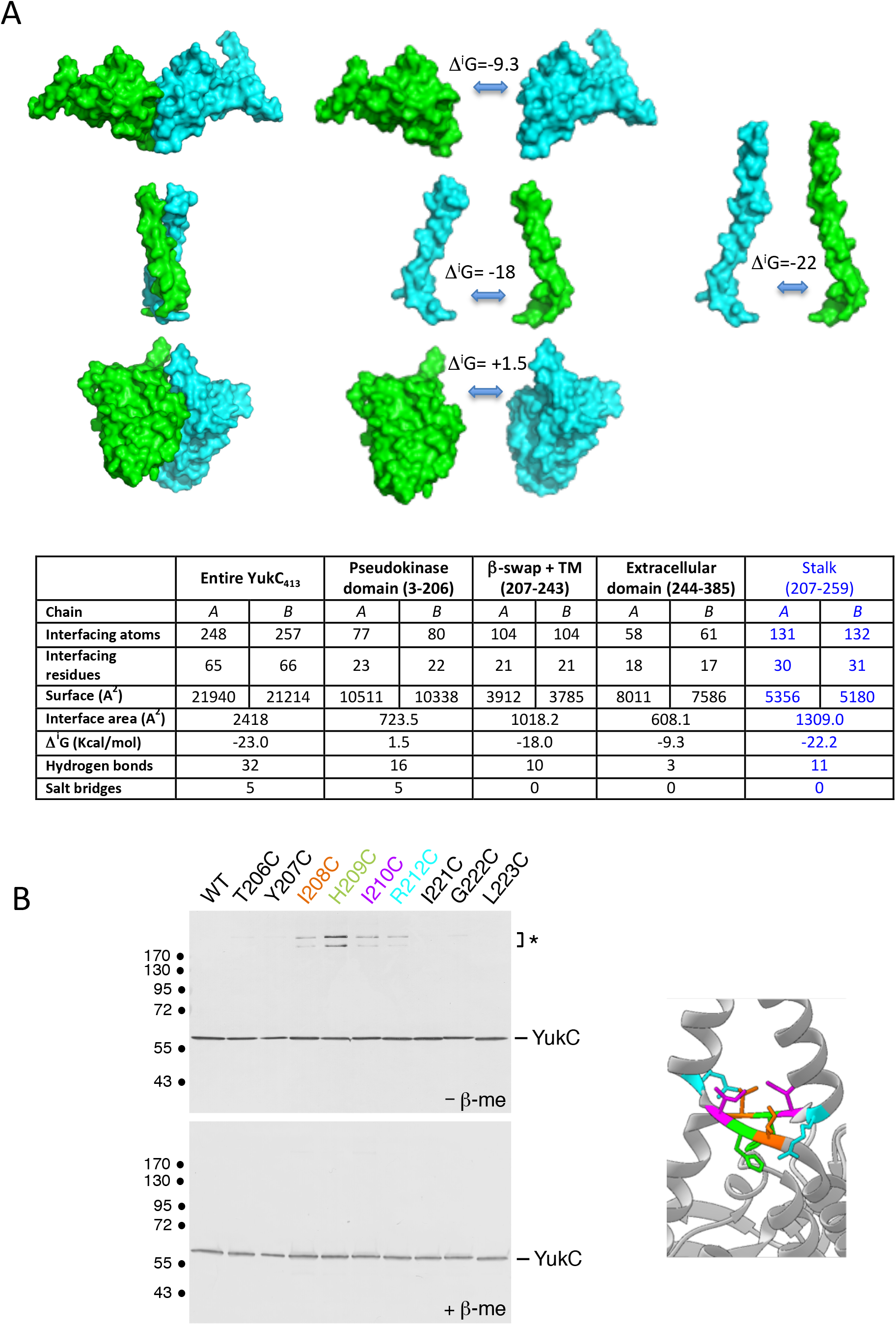
Structural bases of YukC dimer interaction. **A)** Energetics of YukC_413_ monomers interaction. Chain A is rendered in cyan and chain B in green. Δ^i^G indicates the solvation free energy gain upon formation of the interface. Top: schematic of the different contributions to YukC_413_ dimer stabilization. Each YukC domain is shown separately (left and center) and their interactions compared to the stalk including the β-swap, the TM region and a portion of the extracellular domain (right). Bottom: results from dimer interface analysis by PISA. **B)** *In vivo* validation of the YukC_413_ structure by disulphide cross-linking. Left: total extracts of *B. subtilis* cells producing the ‘wild-type’ His6-YukC_413_ protein or the indicated cysteine variants were subjected to SDS-PAGE and immunodetection using the anti-His antibody. Samples were treated (lower panel, -β-me) or not (upper panel, +β-me) with β-mercaptoethanol as reducing agent. The asterisk on right indicates bands corresponding to a disulphide-bond YukC oligomer. The dimerizing residues are indicated with the same colorcode used in the scheme highlighting them on the YukC_413_ structure (left panel).

The YukC stalk spans from the ‘β-swap’ domain up to the beginning of the extracellular domain. While the pseudokinase domain and the extracellular region are highly similar to the structures presented previously [15], the overall YukC organization was completely unpredicted around the transmembrane region presenting the novel β-swap domain and the dimer twisting around the stalk.

In the transmembrane portion of the stalk, each α-helix is tilted at 70° with respect to the *β*-swap domain forming two small cavities, one inside the membrane and one right below (Figure 5-figure supplement 1A). The two helices change their orientation inside the membrane at the level of ‘kink 1’ caused by Proline 231. A second change in helices orientation at the ‘kink 2’ (residues 241-242) brings the two α-helices roughly parallel to each other, forming the top of the stalk in the extracellular domain.

At the base of the stalk, the *β*-swap domain has, to the best of our knowledge, no structural known homologs. Its structure is made of a two-stranded, antiparallel β-sheets running roughly parallel to the membrane plane, where the two monomers cross before regaining a α-helical organization. Interestingly, a cluster of hydrophobic amino acids in the *β*-swap (I^208^, I^210^ and P^211^) forms a hydrophobic patch located below the transmembrane region and capped by the W^214^ residue coming from the stalk (Figure 5-figure supplement 1B). Right in the middle of the *β*-swap, the two H^209^ point their lateral chains in opposite direction with respect to the hydrophobic patch and towards the pseudokinase domain (Figure 5-figure supplement 1B, inset). Before and after the *β*-swap domain, three residues (K^205^, R^212^ and K^213^) define a positively charged region that may interact with the negatively charged polar heads of the lipids (Figure 5-figure supplement 1C).

Because the *β*-swap domain represents a novel structural feature, we sought to verify this structure *in vivo*. To this purpose we performed a cysteine-scanning analysis of the twist region. Residues 206-212 and 221-223 were individually substituted by cysteines, and the formation of disulfide bonds between the two monomers was assessed by denaturing SDS-PAGE in reducing or non-reducing conditions. Intermolecular disulfide bridges were detected in mutants I208C, H209C, I210C and R212C (Figure 5B). The strongest effect was observed for the H^209^ residue substitution, facing each other in the *β*-swap (Cα-Cα distance in YukC_413_ model = 5 Å), while I^208^ and I^210^ presented a weaker effect (Cα-Cα distance = 8.1 and 8.4 Å, respectively). Surprisingly, mutants in R^212^ were also able to form disulfide bonds despite their distance in the crystallographic model (15.6 Å). The formation of a disulfide bond in this case suggests that the dimer may undergo a significant structural change, such as a scissor movement.

Taken individually, the structures of the two monomers in YukC_413_ are basically identical except at the level of the region between the *β*-swap and the end of the pseudokinase domain (Figure 5-figure supplement 2A, red asterisk). Here, the K^205^ residue of monomer A forms a H-bond with the A^80^ of B while, on the other side, the side chain of K^205^ from monomer B is pointing out, making no interactions (Figure 5 -figure supplement 2B). A difference in the orientation of the α-helices 8 between the two monomers produce a 4-Å shift at their top and breaks the symmetry of the YukC dimer (Figure 4 and Figure 5-figure supplement 2B).

## Discussion

In this study, we provide evidence that the *B. subtilis* T7SSb is involved in inter- and intraspecies competition. We then delineate the interaction network between T7SSb subunits, demonstrating a central role for the YukC pseudokinase. Finally, our structure of YukC highlights new features suggesting a connection between YukC flexibility and its role in T7SSb.

### T7SSb dependent bacterial killing in B. subtilis

Bacteria use distinct mechanisms to compete with each other for conquering an ecological niche. In *B. subtilis*, contact-independent competition can occur through production of specialized metabolites such as Bacillaene [43], the lipopeptides plipastatin [44], and surfactin [45]. Surfactin is required for biofilm formation and motility, but also inhibits other bacteria, probably favoring *B. subtilis* fitness in some environments [46, 47]. In addition, *B. subtilis* secretes toxins of the YD-repeat protein family, which inhibits the growth of other *B. subtilis* cells in a contact-dependent manner [48]. Recently, a new class of toxins with a conserved N-terminal (LF)XG motif and a predicted α-helical bundle fold similar to WXG proteins was identified in Gram-positive bacteria [27]. The C-terminal domain of these proteins from *B. subtilis* exhibited RNase activity and were shown to be toxic upon production in *E. coli* [28]. A bioinformatic analysis on several Firmicutes genomes suggested that LXG proteins are secreted through the T7SSb [26]. In agreement with this prediction, the N-terminal domain of the *S. aureus* T7SSb toxin EsaD belongs to the same superfamily than LXG toxins [49]. Recent results obtained in *S. aureus* and *S. intermedius* [26, 49] showed that LXG toxins directly mediate intra-species bacterial killing. However, no evidence was provided so far demonstrating the antibacterial role of T7SSb in non-pathogenic bacteria, including in the model system *B. subtilis*. Here we show that *B. subtilis* is able to cause both intra- and interspecific contact-dependent growth inhibition in a T7SSb-dependent manner. Since six LXG toxins were identified in *B. subtilis*, we reckoned that elimination of one of the toxin-antitoxin pairs would affect bacterial ability to survive in a competition context. Indeed, the *Bs*Δ*yxiD-yxxD* strain is efficiently killed by its *B. subtilis* 168 parental, wild-type, strain, and this activity depends on the presence of the full set of genes encoding the T7SSb (Figure 1A and Figure 1-figure supplement 1A). Therefore, *B. subtilis* 168 produces a functional T7SSb, despite what has been previously proposed on the basis of the lack of detection of the WXG substrate, YukE, in culture supernatants [50]. Discrepancy between these results is likely due to the level of sensitivity of the two methods used to assess T7SSb functionality, a bacterial competition assay being much more sensitive that YukE immuno-detection. Interestingly, the presence of *yukE* is also required for efficient killing, suggesting that (i) YukE plays an active role in the assembly or functioning of the T7SSb, and/or that (ii) YukE interacts with or chaperons secreted toxin(s), or participates to the recruitment of the toxins to the secretion apparatus.

Furthermore, we demonstrated that the *B. subtilis* T7SSb is able to mediate intraspecie competition as well, and in particular against mono-dermal bacteria (Figure 1D and 1E). Interestingly in the time-lapse microscopy movies the disappearance of prey cells starts several hours after the initial contact with attacker cells, both in the intra- and in the interspecies competition (Video 1 and Video 3). This is in accordance with the RNase activity suggested for the LXG toxins in *B. subtilis* [28] which effects may be slower than those of toxins acting on the stability of the bacterial wall. Another possibility is that the T7SSb-dependent killing would be activated by specific factors such as the cell cycle or the density of prey cells. In the future, it will be interesting to further characterize this killing mechanism to understand which factors and environmental cues control T7SSb expression and activation. In the movie of *B. subtilis ΔyukC* interaction with *L. lactis*, the growth of the first one is greatly inhibited, probably by acidification of the media or by the secretion of the bacteriocin nisin by the second one [33]. The lack of inhibition in the presence of a full set of T7SSb encoding genes therefore emphasizes the role of this system in bacterial competition in *B. subtilis* and probably in Firmicutes overall. Di-dermal bacteria proved to be unaffected by T7SSb-dependent competition, independently from their Gram-negative or Gram-positive nature (*E. coli* and *C. glutamicum* respectively) (Figure 1C). Despite a broader investigation is needed to better understand the molecular details of T7SSb-mediated competition (such as the mechanism of toxin transport into the target), our data already suggest that LXG toxins can be transported *via* the T7SSb through the typical monoderm barrier found in Firmicutes, but they seem unable to cross the double membrane of actinobacteria or typical Gram-negative bacteria. In the future, structural information will help understanding how these toxins are delivered into prey cells.

### The T7SSb interaction network in B. subtilis

How the LXG toxins contact and are transported through the T7SSb is still not understood. In *S. intermedius* they were proposed to interact with WXG-like proteins [26] while in *S. aureus*, different effectors (EsaD-EsaE, EsxB, EsxC and EsxD) contact the C-terminus of the coupling protein [14, 21], suggesting that different mechanisms would mediate substrates recognition in the T7SSb. Our BACTH results place YukC as an ideal candidate for interacting with LXG toxins mediating their contact with the rest of the T7SSb (Figure 2C). Moreover, the interaction of YxiD N-terminus with YukE, and the requirement of the latter for bacterial competition support the idea that WXG100 proteins may act as recruiting factors or chaperons for LXG toxins.

The interaction network of T7SSb subunits yielded a complete picture of the molecular interactions within this secretion complex, broadening the understanding of how T7SSbs are organized, and confirming the sparse data available on the *S. aureus* T7SSb (Figure 2A and Figure 2-figure supplement S3).

In particular, the fact that all the T7SSb subunits directly interact with YukC highlights the importance of this subunit in T7SSb (Figure 2A and 2B). Key YukC interactions with: i) the substrate YukE, ii) the multispanning membrane protein YueB and iii) the ATPase YukB were also verified by co-purification and *in vivo* binding experiments (Figure 3).

Our results also show that the large multi-spanning protein of the *Bs*T7SSb system, YueB, interacts directly with the ubiquitin-like subunit YukD. Interestingly, in T7SSa, the multispanning membrane protein EccD has an N-terminal ubiquitin-like domain. Despite the lack of sequence conservation between the multispanning transmembrane proteins in T7SSa and T7SSb, our result suggests a functional correlation between these subunits. However, the ubiquitin-like domain of EccD3 contacts the cytoplasmic portion of the EccC3ATPase in the ESX-3 T7SSa [7, 8], while the *Bs*T7SSb ubiquitin-like subunit YukD does not contact directly the coupling protein. YukC and/or YueB might mediate such interaction in the *Bs*T7SSb.

The interaction of YukC pseudokinase regions with the FHA domains unveils the role of these domains at the N-terminus of the coupling proteins in T7SSb (Figure 3C and 3D). FHA domains usually recognize and bind phospho-threonine residues on different eukaryotic and prokaryotic proteins [17, 51]. However the phosphorylation of the threonine residues on YukC domain would require an eSTK (also known as Hanks-type) kinase, such as PrkC in *B. subtilis [52]*. Since the YukC-YukB/FHA BACTH and co-purification assays were performed in *E. coli*, where to our knowledge no ‘typical’ eSTK is present [53], this suggests that this FHA-pseudokinase interaction is phospho-independent. Indeed, typical residues responsible for the binding of phosphorylated targets are not present neither in the FHAs of YukB [17], nor in those of EssC from *S. aureus* and *G. thermodenitrificans* (PDB 1WV3 and 5FWH, respectively) [12, 13], supporting the phospho-independent nature of the YukB-YukC interaction.

### The YukC_413_ structure and its functional implications

The YukC_413_ crystal structure provides novel information to help understanding the possible mechanism of this subunit in T7SSb complexes. The extracellular domain of YukC does not appear to be directly involved in the multiple interactions observed with the other membrane components of T7SSb. Indeed, our data suggest that YueB-YukC interaction is not mediated by their extracellular regions (Figure 4-figure supplement 2C), while the others *Bs*T7SSb subunits do not have significant extracellular domains. We showed that the C-terminal region of YukC_413_ extends about 40 Å above the cellular membrane (Figure 3-figure supplement 1C) being located inside the peptidoglycan. However the 73 C-terminal residues missing in our construct are predicted to be highly disordered, maybe reaching the external portion of the peptidoglycan. Given its location, this region might be involved in positioning the T7SSb through YukC interaction with the peptidoglycan, as proposed for the *S. aureus* homolog EssB [15].

Spanning from the extracellular domain to the intracellular region, the stalk constitutes an unexpected structural feature producing the YukC monomers rotation around a central axis and stabilizing their dimerization (Figure 5A). It is interesting to note that several proline residues are present along the YukC stalk (Figure 5-figure supplement 2C). It is well known that proline could introduce flexibility in α-helices, in some cases influencing the mechanisms of signal transduction [54–57]. This has been recently proposed for the TM histidine kinase DesK, controlling membrane lipid homeostasis in *B. subtilis* [58]. Interestingly, the distance between key prolines and the orientation of TM helices are striking similar when comparing the model of DesK TM region with the YukC_413_ structure. Given that these prolines were shown to be crucial for DesK signal sensing and transduction [58], it will be interesting to assess their role in the functionality of YukC and T7SSb.

In the crystal packing, two YukC_413_ dimers associate in a head-to-tail orientation, with the pseudokinase domains of one dimer interacting with the extracellular region of the other (Figure 4-figure supplement 1B). Despite these asymmetric contacts, the YukC monomers show structures that mostly overlap (within experimental error) suggesting that the conformation of YukC single domains might not be easily disrupted by contacts with other proteins. However, the monomers differ in the orientation of the region between the last α-helix of the pseudokinase and the *β*-swap (Figure 5-figure supplement 2A). Here, two residues (K^205^ from chain A and A^80^ of chain B) interacting only on one side of the model, participate in the rigid body movement of one pseudokinase domain, which is lifted towards the membrane compared to the other (Figure 5-figure supplement 2B). As discussed above, YukC pseudokinase domains interact directly with the coupling protein (Figure 2, Figure 2-figure supplement 1E, and Figure 3) and maybe with other T7SSb subunits. It will be interesting to analyze the effects of mutations in these two residues (K^205^ and A^80^) on the interaction of the pseudokinase and on T7SSb functionality.

In parallel, the Δ^i^G value of the pseudokinase domains interface suggests a non-stable association of these domains (Figure 5A), which could possibly be modified by the binding of other subunits or substrates in the T7SSb. Moreover, we obtained experimental evidences of a possible lateral flexibility of the *β*-swap using cysteine-scanning analysis. With this experiment we also confirmed the nature of this peculiar structural arrangement by proving that the three central residues of the strands are close enough to form disulfide bonds. Interestingly, whereas in the YukC_413_ model, R^212^ residues are located quite far from each other (15.6 Å) their substitution induces disulfide bridges in YukC (Figure 5B). Therefore an *in vivo* conformational change of this region should occur to allow the two side chains to be in contact. A scissor-like movement of the β-strands at the level of the swap region could bring these residues close to each other at a certain point of the YukC activity. Alternatively, formation of the sulfur bridge might destroy the β-swap. In both cases, our result shows that the pseudokinase domains are flexible enough to form this sulfur bridge. The hydrophobic patch located between the stalk and the TM region (residues I^208^, I^210^, P^211^ and W^215^, Figure 5-figure supplement 1B) could play a role in this mechanism, especially in the case of T7SSb assembly in lipid rafts proposed in [59].

Altogether, our evidences support a model where YukC undergoes dynamic structural changes in its intracellular region that could modulate its interactions with other T7SSb subunits and substrates, including the LXG toxins involved in bacterial competition.

## Conclusion

Taken together our results show that *B. subtilis* T7SSb is a *bona-fide* functional T7SS. The many advantages of this bacterial model (non-pathogenic, highly amenable to genetic manipulation and with a well characterized physiology) combined to the simple organization of its T7SSb (compared to *M. tuberculosis* T7SS or *S. aureus* T7SSb), establish *B. subtilis* T7SS as an important model system. Our study lays the foundations of further investigations that will help understanding the complete assembly and secretion mechanism of these important bacterial secretion systems.

## Materials and methods

### Cloning

Plasmids used in this work are listed in Table S2 (plasmids for BACTH assay), Table S3 (plasmids for protein expression) and Table S4 (plasmids for competition assays). Plasmid propagation and construction were routinely performed with *E. coli* DH5α. Genes of interest were amplified with the Phusion polymerase (Thermo Scientific) using purified *B. subtilis* 168 DNA as template and cloned with standard restriction enzymes-based procedures. In the case of *pRSFduet/yueB.strep* four primers were used to generate annealing over-hangs. *pJM14* (*amyE::*P*hyperspank-gfp, Spectinomycin*) was built in a two-way ligation with a PCR amplified fragment containing the *gfp* gene and an optimized ribosome-binding site (oligonucleotide primers oJM31 and oJM32 and plasmid pKL147 [60] as template) and pDR111 (kind gift from D.Z. Rudner) cut with *Hind*III and *Nhe*I. Inserts and vectors were digested with restriction enzymes (New England Biolabs, NEB). Primers used and generated plasmids are listed in Tables S1 to S4.

### Bacterial strains

Strains used are listed in Table S5. To engineer the *B. subtilis* T7SSb mutant strains used in this study (*Bs*Δ*yukE*, *Bs*Δ*yukD*, *Bs*Δ*yukC*, *Bs*Δ*yukB*, *Bs*Δ*yueB* and *Bs*Δ*yueC*), we initially purchased deleted strains from the Bacillus Genetic Stock Center (BGSC) in the *B. subtilis* 168 *trpC*^-^ genetic background carrying the erythromycin (erm) resistance cassette flanked by Lox sites replacing the wild-type gene (BKE31910 to BKE31850). The respective markerless deletions used for our experiments were obtained by eliminating the erythromycin resistance cassette from the *locus* of the respective T7SSb gene from each *B. subtilis* mutant. To this purpose, each strain was transformed with the thermosensitive vector pDR244 encoding for the Cre recombinase [61]. The strains lacking erythromycin resistance were further tested by PCR and pDR244 was eliminated by growing the cells at 42 °C on LB-agar plates.

As ‘attacker’ strains in the bacterial competition, the *B. subtilis 168 CA* wild type or the markerless mutants described above were used. As a ‘prey’ strain for the intra-specie competition we used *B. subtilis* 168 CA lacking the toxin-antitoxin pair YxiD-YxxD. The *Bs*Δ*yxiD-yxxD* strain was made by transforming *B. subtilis* 168 CA with the pJM14 integrative vector carrying the *yxiD-yxxD* flanking sites inserted using Gibson assembly kit from NEB. For inter-species competition *L. lactis subsp. cremoris* (strain NZ9000) was used, carrying the plasmid pGTP_FZ_301 conferring chloramphenicol resistance [62]. To test competition against Gram-negative bacteria, the *E. coli* K-12 W3110 was used, carrying the plasmid *pUA66-rrnB* for constitutive Green Fluorescent Protein (GFP) expression [63]. Competition against diderm Gram-positive bacteria was performed using a strain of *C. glutamicum* carrying the plasmid pUMS-6 [64] with the ‘mScarlet’ fluorescent gene [65] and conferring Kanamycin resistance.

### B. subtilis genome extraction

A fresh single colony was inoculated in 2mL LB and grown until mid to late exponential grown (λ = 600 nm (*A*_600_) ~ 0.8) and the corresponded cells pellet was resuspended in 200 μL of lysis buffer (20mM Tris-HCl, pH 7.5, 2 mM EDTA, 20 mg/mL lysozyme, 1.2 % Triton x100). The bacterial solution was incubated 30 minutes at 37 °C. Subsequent steps followed the ‘DNeasy Blood & Tissue Kits’ protocol (Quiagen – Ref 69504).

### Transformation of B. subtilis

2 mL of LB were inoculated with a *B. subtilis* single fresh colony and grow overnight at 28°C with 180 rpm shaking. Cells were diluted 1:100 in 5 mL of SpI media (2g/L (NH4)_2_SO_4_, 14 g/L K_2_HPO_4_, 6 g/L KH_2_PO_4_, 1 g/L Na-Citrate • 2H_2_O, 0.2 g/L MgSO_4__7H_2_O, 0.5 % glucose, 0.1 % yeast extract, 0.02 % casaminoacids) and grown at 37 °C under agitation at 180 rpm. 0.5 mL of the late exponentially growing cells was added to 4.5 mL of SpII medium (SpI supplemented with 0.5 mM CaCl_2_, 2.5 mM MgCl_2_) and grown 90 min at 37 °C with 100 rpm shaking. The culture was supplemented with 1 mM ethylene glycol-bis(β-aminoethyl ether)- *N,N,N’,N’*-tetraacetic acid (EGTA) and incubated for additional 10 min. 1 to 10 μg of purified DNA was added to 300 μL of *B. subtilis* competent cells and incubated at 30 °C for 90 min with 100 rpm shaking. The culture was spread on LB solid media supplemented with 2 μL/mL Erythromycin.

### Bacterial competition assay

Single fresh colonies from glycerol stocks (strains listed in Table S5) were grown on LB solid media (agar 1.5 %) overnight at 30 °C before being inoculated in 3 mL of LB liquid media supplemented with Spectinomycin (Spec) 100 μg/mL for *B. subtilis* prey cells. Cells were diluted 1:100 in 4 mL of SIM media (Secretion Induced Media: M9 supplemented with 10 % LB, 2.4 % glycerol, 0.4 % glucose, 10 μg/mL thiamine, 75 μg/mL casaminoacids, 1 mM MgSO_4_, 0.1 mM CaCl_2_, 50 μM Fe_3_Cl and 100 μM citrate) without antibiotic. The *B. subtilis* ‘prey’ strain lacking YxiD and YxxD were induced by the addition of 200 μM isopropyl β-D-1-thiogalactopyranoside (IPTG) when cultures reached λ=600 nm (*A*_600_) of 0.2. Two mL of each bacterial culture were pelleted at *A*_600_=0.8 by centrifugation at 8,000 × *g* for 5 min and resuspended in SIM media to an *A_600_* of 10. Predator and prey cells were mixed in a 5:1 ratio, and adjusted to 100 μL with SIM media. 12.5 μL of each bacterial mixture were spotted on SIM solid media (1.5 % agar, supplemented with IPTG 200 μM) in duplicate and the plates were incubated overnight at 30°C. The competition plates were imaged by visualizing GFP fluorescence in a BioRad ChemiDoc system, using the program ‘Alexa535’. Bacteria from each spot were resuspended in 1 mL of LB liquid media and kept on ice. The *A*_600_ was adjusted to 0.5 and 200 μL of each bacterial solutions was placed in a black 96-wells plate, in triplicates. Three wells were filled with sterile LB as negative control. The 96-plate was analyzed using a TECAN microplate reader with excitation at λ=485 nm and emission at λ=530 nm. 100 μL of 10^−3^ to 10^−6^ serial dilutions were plated on LB solid media supplemented with Sp 100 μg/mL. The plates were incubated overnight at 37 °C and the number of surviving *B. subtilis* prey cells were counted. Six spots were analyzed from 3 independent experiments for each prey-predator mix obtaining survival cell data with calculated P values < 0.05.

### Live microscopy

*B. subtilis* and *L. lactis* cultures grown as previously described for competition tests, were applied on SIM media-based 2% agarose pads. Live imaging was performed using a Zeiss Axio Observer Z1 microscope fitted with an Orca Flash 4 V2 sCMOS camera (Hamamatsu) and a Pln-Apo 63X/1.4 oil Ph3 objective (Zeiss) and analyzed using the software Fiji [66]. For *B. subtilis* 168 or *Bs*Δ*yukC vs BsΔyxiD-yxxD*, frames were collected every 20 min during overnight growth at 30 °C, using phase-contrast and epifluorescence microscopy (585 nm, 40 ms exposure time). For *B. subtilis* 168 or *Bs*Δ*yukC vs L. lactis* competition movies, frames were collected only in phase-contrast every 15 min under the same experimental conditions.

### SPP1 infection assay

A fresh *B. subtilis* colony was inoculated in 3 mL of LB liquid media and grew over night at 37 °C, 180 rpm. The culture was diluted 1:100 in 3 mL of LB and once it approached the exponential phase (λ = 600 nm (*A*_600_) 0.4 – 0.6) 250 μL were mixed with 5 mL of LB top-agar (0.6 % agar) supplemented with 10 mM CaCl_2_. The mixture was quickly poured onto LB solid media (1.5 % agar) and waited for its solidification. 10 μL of SPP1 serially diluted solutions were spotted on the top-agar surface and the plate was incubated at 37 °C over night. The SPP1 stock solution was kindly provided by Dr. Paulo Tavares.

### Bacterial Two-Hybrid Assay

The Bacterial Two Hybrid (BACTH) assay was performed according to the manufacturer instructions (Euromedex). Plasmids used for the BACTH assays are listed in Table S2. Briefly, 50 μL of *E. coli* BTH101 cells [37] were co-transformed with 20 μg of pKT25/*X* or pKNT25/*X* and 20 μg of pUT18/*X* or pUT18C/*X* where *X* indicates the gene of interest (see Table S2). As a negative control, pKT25 and PUT18C were used, while the vectors carrying the *zip* gene were used as a positive control. Freshly transformed colonies were picked, growth in 5 mL of LB media supplemented with the selective antibiotic until *A*_600_ = 0.4. 1μL was then spotted on LB media supplemented with 40 μg/mL of 5-bromo-4-chloro-3-indolyl-β-D-galactoside (X-gal) (Euromedex), ampicillin (Amp, 100 μg/mL), kanamycin (Kan, 50 μg/mL) and 0.5 mM IPTG. Alternatively, carbenicillin (Carb) at 50-100 μg/mL was used instead of ampicillin because of its higher stability compared to ampicillin. To increase the amount of colonies tested, a variation of this protocol was developed, which was optimized to give the same results after 72 hours of incubation at 25 °C. Briefly, after double transformation, BTH101 cells were picked and resuspended in 2 μL of LB. Then 1 μL of this mixture was directly spotted on LB solid media supplemented with Carb 50 μg/mL, Kan 50 μg/mL, 5 mM IPTG and 40 μg/mL X-gal. Each transformation was performed at least twice, and 3-6 colonies *per* transformation were tested for each interaction. Therefore each BACTH assay reflects the results of minimum 6 different colonies. The plates were incubated at 25 °C for 72 hours, followed by an additional incubation time at 30 °C for 24 hours when necessary until saturation of the negative control. The X-gal (Euromedex) was freshly prepared before each experiment, by resuspension in dimethylformamide to a stock solution of 40 mg/mL.

### Co-purification of YukC and T7SSb components

#### YukC.his-YueB.strep co-purification

6 L of TB Terrific Broth (TB) supplemented with Carb 50 μg/mL, Kan 50 μg/mL and 0.8 % glycerol were inoculated with *E. coli* C43(DE3) [67] co-transformed with *pET15b/yukC.his* and *pRSF/yueB.strep*. YukC.his and YueB.strep were induced as described for YukC.strep purification (see below). The *E. coli* cells were resuspended in 200 mL of cold Lysis buffer (50 mM HEPES pH 8, 300 mM NaCl, 1 mM EDTA, 5 % glycerol, 4 tablets of protease inhibitors, Roche). The cells were lysed and the membrane fraction was collected as described in the YukC purification protocol below. The membrane fraction was resuspended in 20 mL of cold solubilization buffer (25 mM HEPES, pH 8, 300 mM NaCl, 2.5 % glycerol, 1 mM TCEP, 1 tablet of protease inhibitors). The membrane proteins solubilization was achieved by the addition of 0.5 % Lauryldimethylamine-N-Oxide (LDAO) followed by 40 min incubation under moderate shaking at 4 °C. For the affinity chromatography step, the sample was first loaded into the StrepTrap column (GE Healthcare) and washed with 50 mL of purification buffer (25 mM HEPES, pH 8, 175 mM NaCl, 0.02 % LDAO). The StrepTrap was washed with purification buffer containing 0.004 % Lauryl Maltose Neopentyl Glycol. 2,2-didecylpropane-1,3-bis-β-D-maltopyranoside (LMNG) final instead of LDAO.

Proteins were eluted directly into a 1 mL HisTrap column (GE Healthcare) using purification buffer supplemented with Desthiobiotin 2.5 mM (IBA Solutions For Life Science™). The HisTrap column was then washed with 20 mL of purification buffer supplemented with 40 mM Imidazole and the proteins were eluted with the same buffer supplemented with 0.5 M Imidazole. The eluted proteins were separated on SDS-PAGE and detected by immunoblot.

#### YukC.strep – YukE.his co-purification

2 L of LB supplemented with 50 μg/mL Carb and 50 μg/mL Kan was inoculated with *E. coli* BL21(DE3) co-transformed with *pRSF/yukC.strep* and *pET15b/yukE.his*. YukC_*413*_.strep and YukE.his were induced and cells grown as described below for YukC.strep purification. The YukC.strep-YukE complex was isolated from the membrane fraction applying the protocol used for the YukC.strep purification (see below), except for the presence of 1 % v/v Triton-X 100 instead of 1 % Cymal6. For the affinity chromatography step, the sample was first loaded into the StrepTrap column (GE Healthcare), washed with 50 mL of purification buffer (25 mM HEPES, pH 8, 175 mM NaCl, 0.03 % Triton-X 100) and proteins were eluted directly into a 1 mL HisTrap column (GE Healthcare) using purification buffer supplemented with Desthiobiotin 2.5 mM (IBA Solutions For Life Science™). The HisTrap column was then washed with 20 mL of purification buffer supplemented with 40 mM Imidazole and the proteins were eluted with the same buffer supplemented with 0.5 M Imidazole. The eluted fractions were analyzed on SDS-PAGE. The YukC and YukE identities were confirmed by LC-MS/MS.

#### Co-purification of YukC.strep and YukB_Δ256_ (FHA domains)

2 L of LB supplemented with Carb 50 μg/mL and Spec 50 μg/mL were inoculated with *E. coli* BL21(DE3) *[68]* co-transformed with *pRSF/yukC.strep* and *pCDF/yukB_Δ256B_.his*. In this case, the buffer solutions were supplemented with 0.5 mM TCEP. The YukC_Δ217_-YukB_Δ256_ complex was purified by double affinity chromatography as previously described for the YukC.strep-YukE.his complex. The eluted protein were separated on SDS-PAGE and detected by immunoblot.

#### Co-purification of YukC_Δ217_ (pseudokinase domain) and YukB_Δ256_ (FHA domains)

2 L of LB supplemented with Carb 50 μg/mL and Spec 50 μg/mL were inoculated with *E. coli* BL21(DE3) co-transformed with *pRSF/yukC_Δ217_.strep* and *pCDF/yukB_Δ256B_.his*. In this case, the buffer solutions were supplemented with 0.5 mM TCEP. The YukC YukC_Δ217_-YukB_Δ256_ complex was purified from the soluble fraction of the lysate by double affinity chromatography as previously described for the YukC.strep-YukE.his complex. The eluted protein were separated on SDS-PAGE and detected by immunoblot.

### Immunoblot

The proteins separated on SDS-PAGE were transferred on a PVDF membrane (0.45 μM, Merk) previously activated by ethanol and pre-equilibrated with transfer buffer (25 mM Tris, 192 mM glycine, 0.1% SDS, 20% ethanol) with a Trans-Blot Turbo system (BioRad). The PVDF membrane was blocked for 1 h in PBS supplemented with 0.1 % Tween-80 from Sigma Aldrich (‘PBS-Tween’) and 4 % of milk powder. The primary antibodies (diluted 1:1000 for the anti-Strep, and 1:2000 for the anti-His6) were incubated in the same buffer overnight at 4 °C under gentle shaking. Anti-Strep antibodies were purchased from IBA Solutions For Life Science™, while the anti-His6 was from Sigma Aldrich. The PVDF membrane was washed three times for 10 min in PBS-Tween. The secondary antibody antimouse HRP-conjugated (Sigma Aldrich) was diluted 1:5000 in PBS-Tween. After 1 h incubation, the PDVF membrane was washed 3 times for 10 min in PBS-Tween. The blots were developed with ECL (Amersham Biosystem) and revealed in a ChemiDoc system (BioRad).

### *YukC.strep and* YukC_413_ *purification*

A single fresh colony of *E. coli* BL21(DE3) transformed with pRSF/yukC.strep or pRSF/yukC_413_.strep was inoculated in 100 mL of LB liquid media supplemented with Kan 50 μL/mL. After overnight growth at 37 °C overnight shaking at 180 rpm, the bacterial preculture was diluted 1:50 in 4 L of LB media supplemented with Kanamycin (50 μL/mL) and this large-scale culture was incubated at 37 °C, 180 rpm until λ = 600 nm (*A*_600_) 0.8, when YukC.strep production was induced by the addition of 0.5 mM IPTG while the incubation temperature was moved to 16 °C. The culture was incubated overnight shaking at 180 rpm. Cells were then collected by centrifugation at 6000 × *g* for 20 min at 4°C. During the following purifications steps the samples was kept on ice unless otherwise specified. The *E. coli* cells were resuspended in 100 mL of cold lysis buffer (50 mM HEPES pH 8, 175 mM NaCl, 5 % glycerol, 1 mM EDTA and 2 tablets of protease inhibitors by Roche). The bacterial solution was then incubated at room temperature (RT) for 20 min under moderate shaking in the presence of 0.5 mg/mL lysozyme (Sigma-Aldrich) at and additional 10 min after addition of 0.1 mg/mL DNase, 5 mM MgCl_2_ and 1 mM TCEP. The bacterial cells were lysed through 3 cycles into an Emulsiflex (Avestin, US) keeping the pressure between 15 000 and 20 000 psi. In order to remove the unbroken cells, the lysate was centrifuged for 15 min at 12000 × *g*. The membrane fraction was collected by ultracentrifugation in a Ti45 rotor (Beckmann) running at 80000 × *g* for 1 h. The soluble fraction was discarded and the membrane fraction resuspended in 20 mL of cold solubilization buffer (25 mM HEPES, pH 8, 175 mM NaCl, 2.5 % glycerol, 1 tablet of protease inhibitors). 20 mL of solubilization solution supplemented with the 2 % 6-Cyclohexyl-1-Hexyl-β-D-Maltoside (Cymal6, Anatrace) was added gently to the membrane solution and the sample incubated at RT for 30 min under moderate shaking. Unsolubilized material was removed by ultracentrifugation using the rotor SW32 (Beckmann) at 50000 × *g* for 45 min. The cleared membrane fraction was loaded with a flow of 0.8 mL/min on a pre-equilibrated 5 mL StrepTrap column (GE healthcare) mounted on as AktaPurifier system (GE healthcare) in the presence of purification buffer (25 mM HEPES, pH 8, 175 mM NaCl, 0.04 % Cymal6). The affinity column was washed using 50 mL of purification buffer. YukC was eluted using the purification buffer supplement with 2.5 mM Desthiobiotin (Sigma). The protein composition of the eluted fractions and the intermediated purification steps were evaluated by SDS-PAGE. 5 mL of YukC solution was injected on a Superdex 200 PG 16/600 column to further eliminate contaminant proteins. The yield of protein was approximately 5 mg *per* L bacterial culture.

### YukC crystallization and structure determination

Crystals of YukC_413_.strep were obtained by hanging-drop vapor diffusion in 0.4 μL with a 1:1 ratio between the protein solution (20 mg/mL YukC_413_, 25 mM HEPES, pH 8, 175 mM NaCl, 0.04 % Cymal6xs and the crystallization solution (0.1 M MgCl_2_, HEPES 50 mM, pH 7.5, 15 %w/v PEG 2K). 200 nl drops were dispensed by a Mosquito Crystal robot (ttp labtech) at 4°C and crystals appeared within three days. The crystals were frozen in liquid nitrogen with a 50% paratone N (Hampton Research), 50% paraffin oil solution as the cryo-protectant. A 2.6-Å resolution dataset was collected on beamline ID30A-3 at the European Synchrotron Radiation Facility (ESRF) in Grenoble, France. Data were processed with autoPROC [69] and the structure (PDB ID: 6Z0F) was solved by a two-step molecular replacement approach with PHASER [70]. Rms deviations were calculated with Molprobity [71] within the phenix crystallographic software suite.

A first molecular replacement search was performed using the coordinates of the dimeric extracellular portion of *G. thermodenitrificans* EssB (PDB 2YNQ) as the probe, followed by a new search using the coordinates of the pseudokinase domain from the same protein (PDB 4ANO). Once a solution including both the intracellular and extracellular domains was identified, the missing part of the molecule, including the transmembrane domain, was modeled combining automated tracing with PHENIX AutoBuild [72] and manual rebuilding with Coot [73]. The overall model was then subject to alternating cycles of inspection with coot and refinement with BUSTER [74].

### Analysis of the YukC_413_ dimer interface

Coordinates of YukC_413_ were analyzed with the PISA server (http://www.ebi.ac.uk/msd-srv/prot_int/cgi-bin/piserver [42]) to evaluate dimerization surfaces (A^2^) for different regions of YukC_413_ as well as their corresponding ΔG^i^s, and to identify residues involved in H-bonds or salt bridges.

### Analysis of the β-swap domain

#### Cysteine substitution construction

Codon substitutions were introduced into the vector pBE-S/*yukC*his by site-directed mutagenesis using complementary pairs of oligonucleotides bearing the desired substitution (listed among others in Table S1, synthesized by Sigma-Aldrich) and the Pfu Turbo DNA polymerase (Agilent technologies). All constructs have been verified by DNA sequencing (Eurofins genomics).

#### Cysteine cross-linking assay

Overnight cultures of *B. subtilis* cells producing the wildtype 6×His-tagged YukC or its cysteine derivatives were diluted to a *A*_600_ of 0.1 in LB. Once the cultures reached *A*_600_ = 0.8, N-ethyl maleimide (Sigma-Aldrich) was added at the final concentration of 5 mM to block all free thiol groups. After 20 min of incubation at 37 °C, cells were harvested, resuspended in lysis buffer (20 mM Tris-HCl pH7.5, 10 mM EDTA, 10 mM MgCl_2_, 1 mg/mL lysozyme, 100 μg/mL DNase) supplemented with protease inhibitors (Complete, Roche), frozen for 15 min at −80°C, and thaw at 37°C before addition of Laemmli loading buffer supplemented or not with β-mercaptoethanol 8 % and boiled for 5 min. Total extracts were run on 10%-acrylamide SDS-PAGE, and transferred onto nitrocellulose. YukC and YukC multimers were detected with anti-histidine-specific antibody (clone AD1.1.10, Biorad).

#### Comparison of YukC_413_ structure

We searched for structures similar to YukC_413_ using the web tool DALI [39] obtaining the rmsd, the values for sequence identity and Z-score. The selected protein sequences were aligned with the web tool T-COFFEE [75] and the structures were super-imposed with the software UCSF Chimera [76].

#### Nano differential scanning fluorimetry (nanoDSF)

Thermal denaturation was performed on the Prometheus NT.48 (Nanotemper) from 20 to 95°C with 80 % excitation laser power and 2 °C/min of heating rate. The tryptophan fluorescence emission was monitored at 330 nm and 350 nm as a function of increasing temperature. YukC and PknB were diluted to a concentration of 1 mg/mL in presence or in the absence of 1 mM ATP in 25 mM HEPES, pH 8, 175 mM NaCl, 2 mM MgCl_2_, 0.04 % Cymal6. The sample was filled into the capillaries and the emission at 350 nm and 330 nm was measured. The intrinsic fluorescence signal expressed by the 350nm/330nm emission ratio was plotted as a function of temperature. Three replicates were performed for each protein. The average of the replicates was calculated and plotted in Figure 4-figure supplement 4.

## Supporting information

Video 1. BsWT vs BsPrey

Video 2. BsDeltayukC vs BsPrey

Video 3. BsWT vs L. lactis

Video 4.BsDeltayukC vs L. lactis

## Acknowledgments

We are thankful to A. Haouz, head of the Crystallography platform (Institut Pasteur, Paris), for his support in YukC crystallization. We acknowledge S. Brûlé and B. Raynal from the Molecular Biophysics platform of the Institut Pasteur (Paris) for performing nanoDSF experiments and analytical ultracentrifugation (not inserted in the current study). We acknowledge the European Synchrotron Radiation Facility for provision of synchrotron radiation facilities and we would like to thank Gordon Leonard for assistance in using the beamline ID30A-3. We also thank the Synchrotron SOLEIL and its staff for assistance in using the beamline to test several YukC crystals. We are grateful to P. Tavares (I2BC, Université Paris-Sud, Université Paris-Saclay) for its kind gift of purified SPP1 phage. We acknowledge the Pasteur proteomics platform at the Institut Pasteur (Paris) for performing the MS analysis of YukE. We are grateful to A. Wehenkel, O. Francetic and S. Dramsi (IP, Paris) for the many helpful discussions and to N. Minc (IJM, Paris) for his help with data and imaging analyses. We thank G. Pehau-Arnaudet of the ‘Unité Technologie et service Biolmagerie Ultrastructurale’ at the Institut Pasteur (Paris) for his support in the initial electron microscopy investigation on YukC. We also thank L. Catorie and M. Casiraghi (IBPC, Paris) for the insertion of YukC in nanodiscs, even if the results could unfortunately not be inserted in the current study. This work received financial support from the Institut Pasteur (Paris). M.T. was supported by a scholarship from the Pasteur-Paris University (PPU) International PhD Program.

---

**Figure 1-figure supplement 1.**
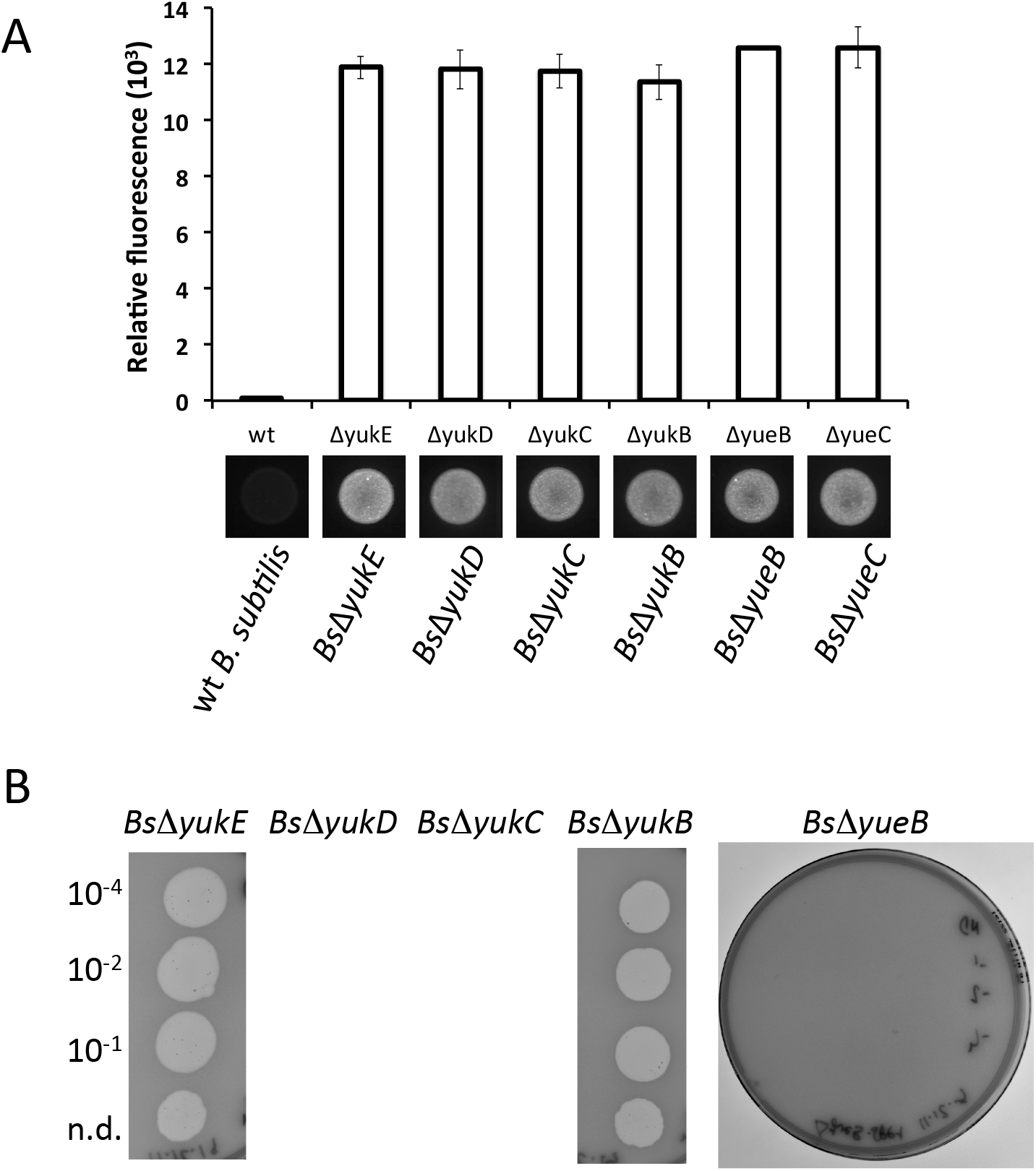
Competition assays controls. **A)** GFP emission quantification of the correspondent experiments reported in Figure 1A. The bacterial spots after overnight competition were resuspended in minimal media, normalized to the bacterial OD_600_, and their fluorescence measured at 530 nm (excitation 485 nm). The pictures of spots fluorescence for the corresponding samples are reported below. **B)** Effect of T7SSb gene deletions on the ability of SPP1 phage to lysate *B. subtilis*. Phage-induced bacterial lysis, mediated by the protein YueB, is observed in each deletion of the first four genes of the operons (namely *Bs*Δ*yukE*, *Bs*Δ*yukD*, *Bs*Δ*yukC* and *Bs*Δ*yukB*), while deletions in the *yueB* gene causes resistance of *B. subtilis* to SPP1 lysis.

**Figure 2-figure supplement 1.**
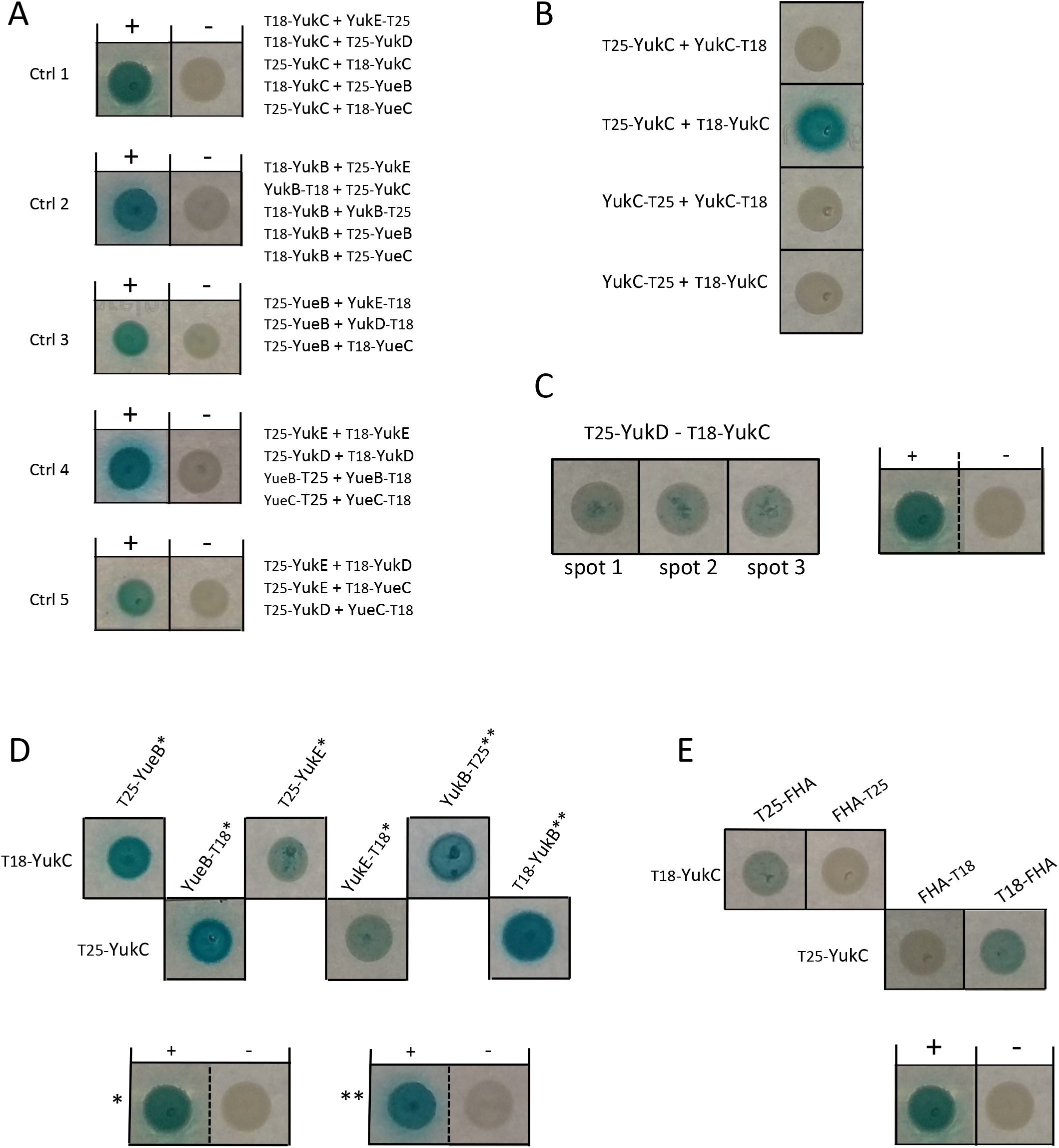
*Bs*T7SSb BACTH assay. **A)** Series of BACTH controls for each sets of experiments shown in Figure 2A. The list of the correspondent experiments is reported on the right side of the control image. **B)** BACTH results on YukC oligomerization confirming its predicted topology with the N-terminus inside the cell and the C-terminus outside. **C)** Three different YukC-YukD interaction spots are reported showing the consistent positive interaction compared to the controls. **D)** The interaction of YukC with YueB, YukE and YukB is independent from the side of the T25- or the T18-tags. **E)** BACTH assay between YukC and the N-terminal domain of YukBΔ256 including the two FHA regions (‘FHA’).

**Figure 2-figure supplement 2.**
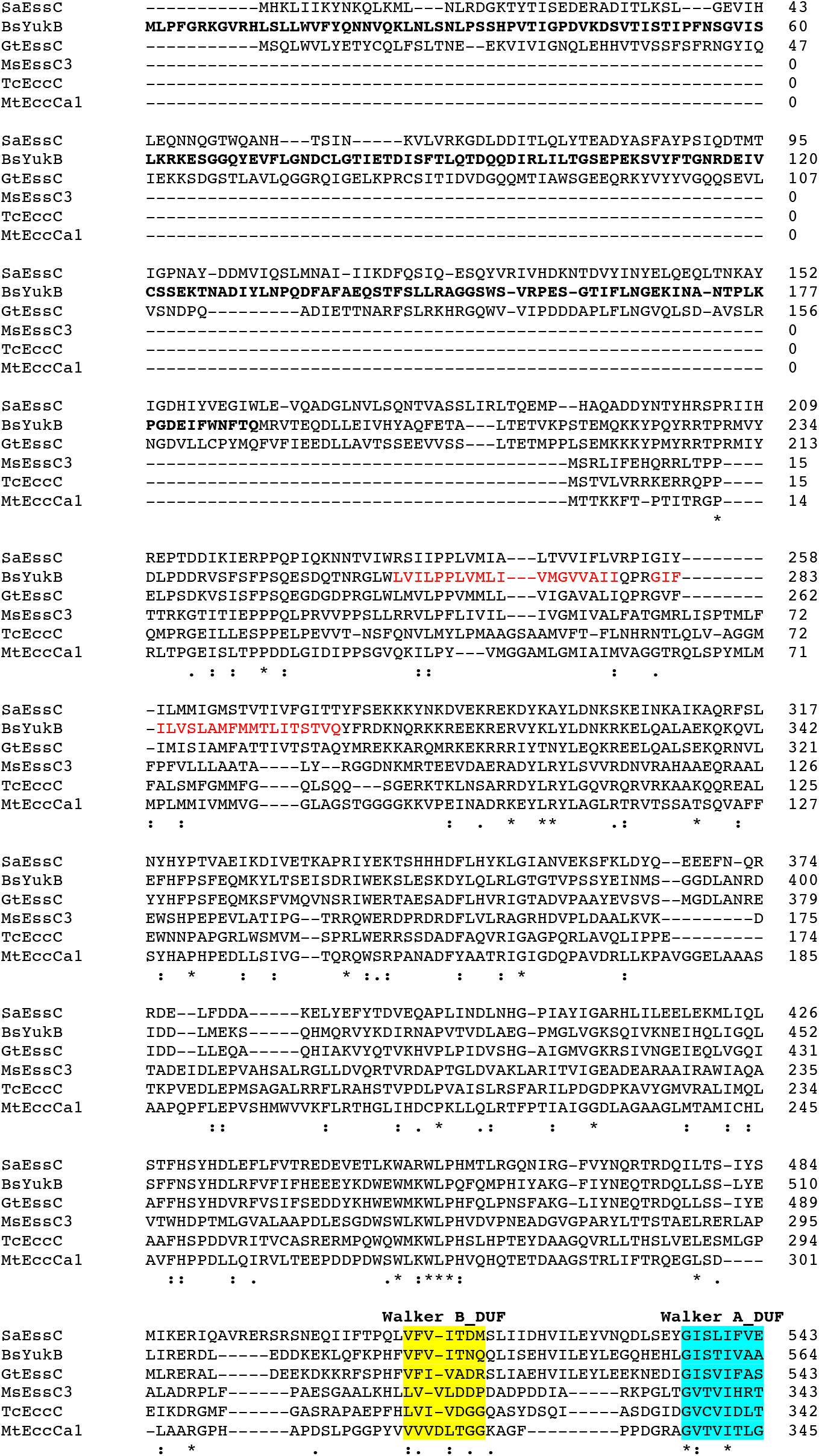
Sequence alignment between YukB and selected homologs from T7SS and T7SSb. Sequences of six coupling proteins were aligned from the beginning to the ATPase region of the DUF domain using the Clustal Omega (1.2.4) program. The following protein were analysed: EssC from *S. aureus* USA300 (SaEssC), YukB from *B. subtilis* 168 (BsYukB), EssC from *G. thermodenitrificans* (GtEssC), all from T7SSb, and the following coupling proteins from T7SSa: EccC from *M. smegmatis* ESX-3 (MsEccC3), the *T. curvata* (TcEccC), and EccCa from *M. tuberculosis* ESX-1 (MtEccCa1). The FHA domains are depicted in bold characters, the TM regions in red and the Walker A and B regions of the DUF ATPase are highlighted in cyan and yellow, respectively. Fully conserved residues are indicated by asterisks (*), colons (:) indicate conservation between groups of strongly similar properties and semicolon (;) conservation between groups of weakly similar properties.

**Figure 2-figure supplement 3.**
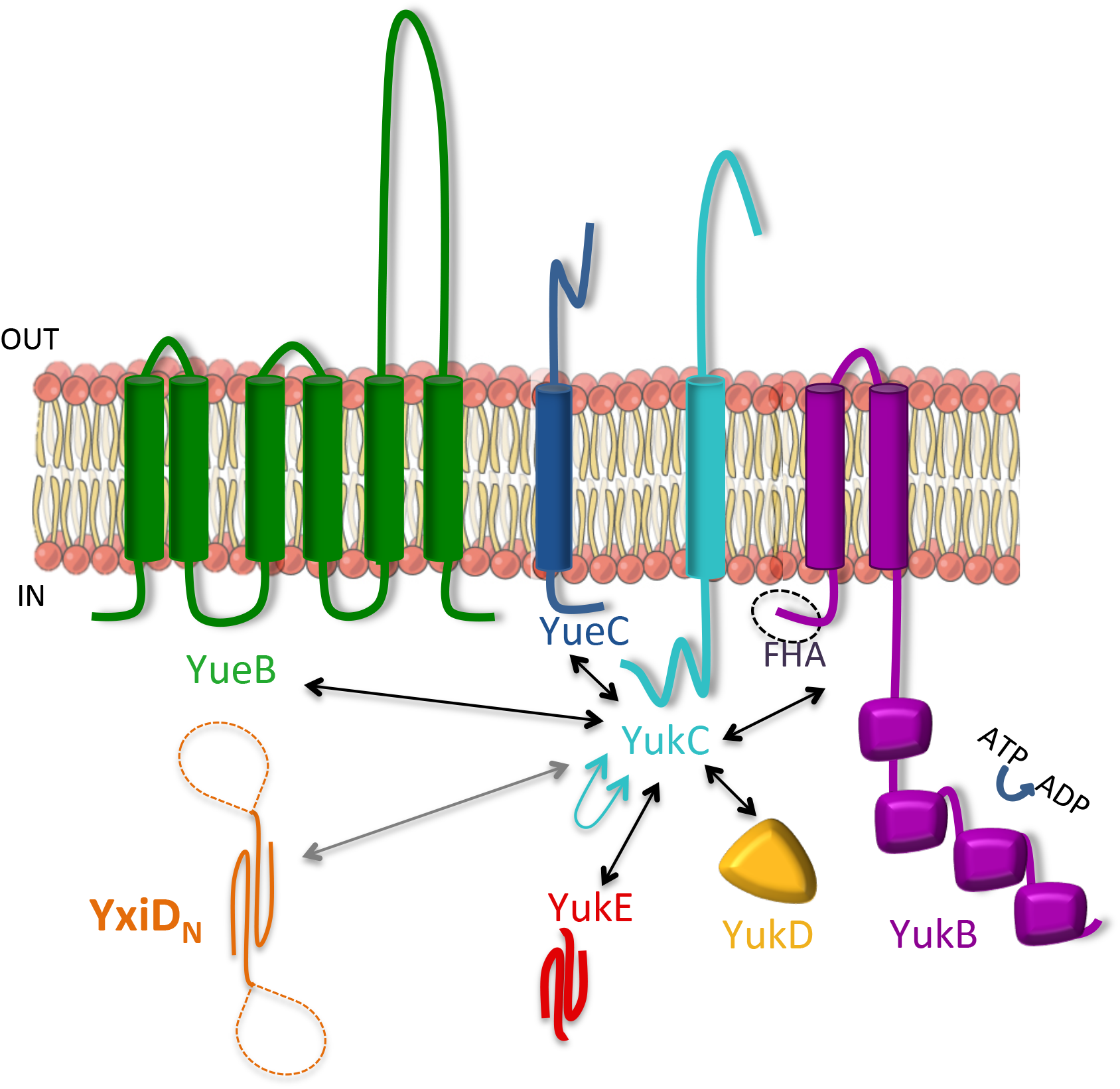
Schematic of YukC interactions with the T7SSb components and with YxiD_N_. The T7SSb components are represented accordingly to their predicted topology. YukC and its self-interactions are represented in cyan. Black arrows represent interactions between YukC and T7SSb subunits. Interactions of the N-terminal domain of YxiD toxin (YxiD_N_, in orange) are indicated in grey.

**Figure 4-figure supplement 1.**
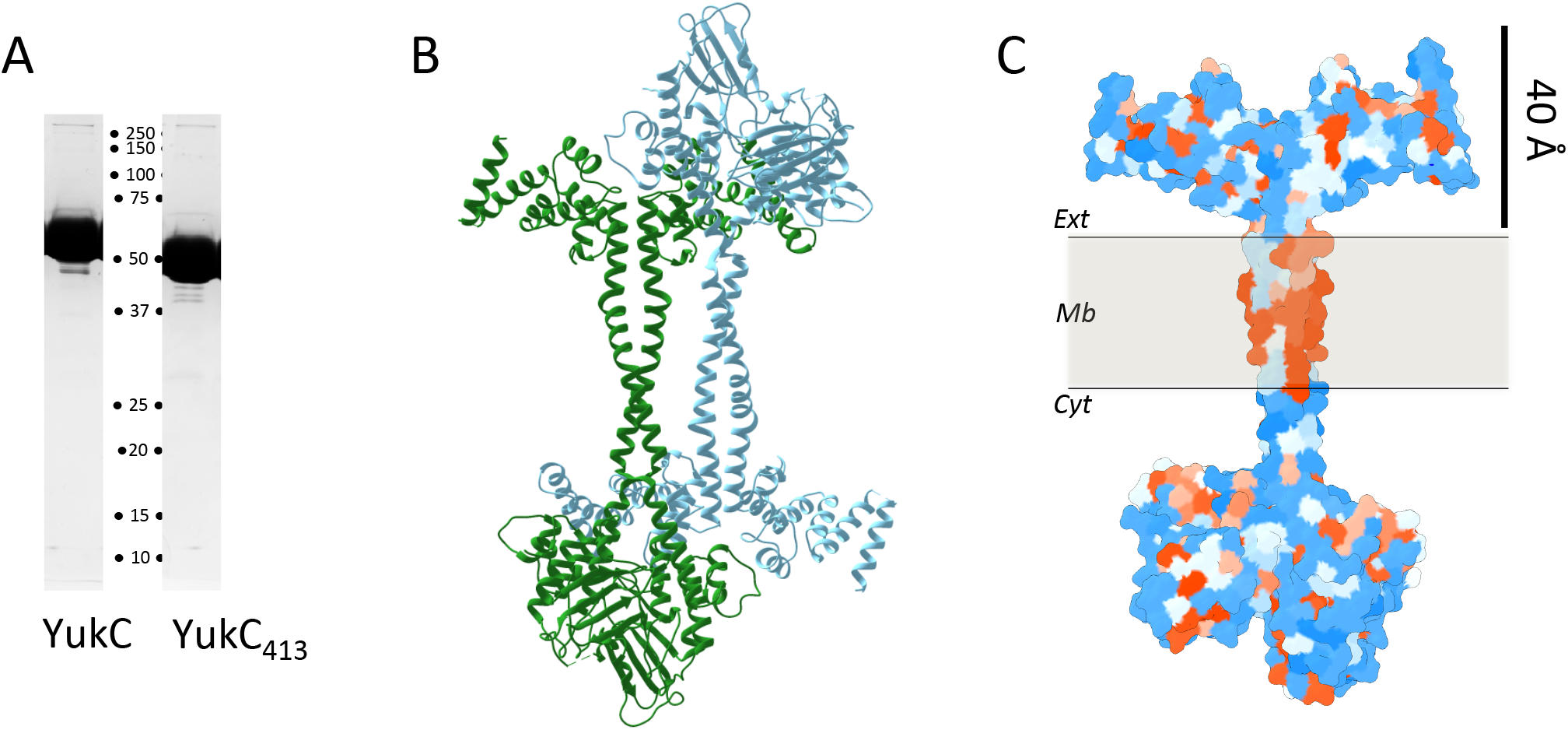
Crystallization of YukC_413_. **A)** SDS-PAGE migration of purified YukC (left) and YukC_413_ (right). **B)**. Head-to-tail arrangement of YukC_413_ dimers in the crystal packing. **C)** Atomic model of YukC_413_ superimposed with the hydrophobicity surface as calculated by the Chimera software (red: hydrophobic, blue: hydrophilic).

**Figure 4-figure supplement 2.**
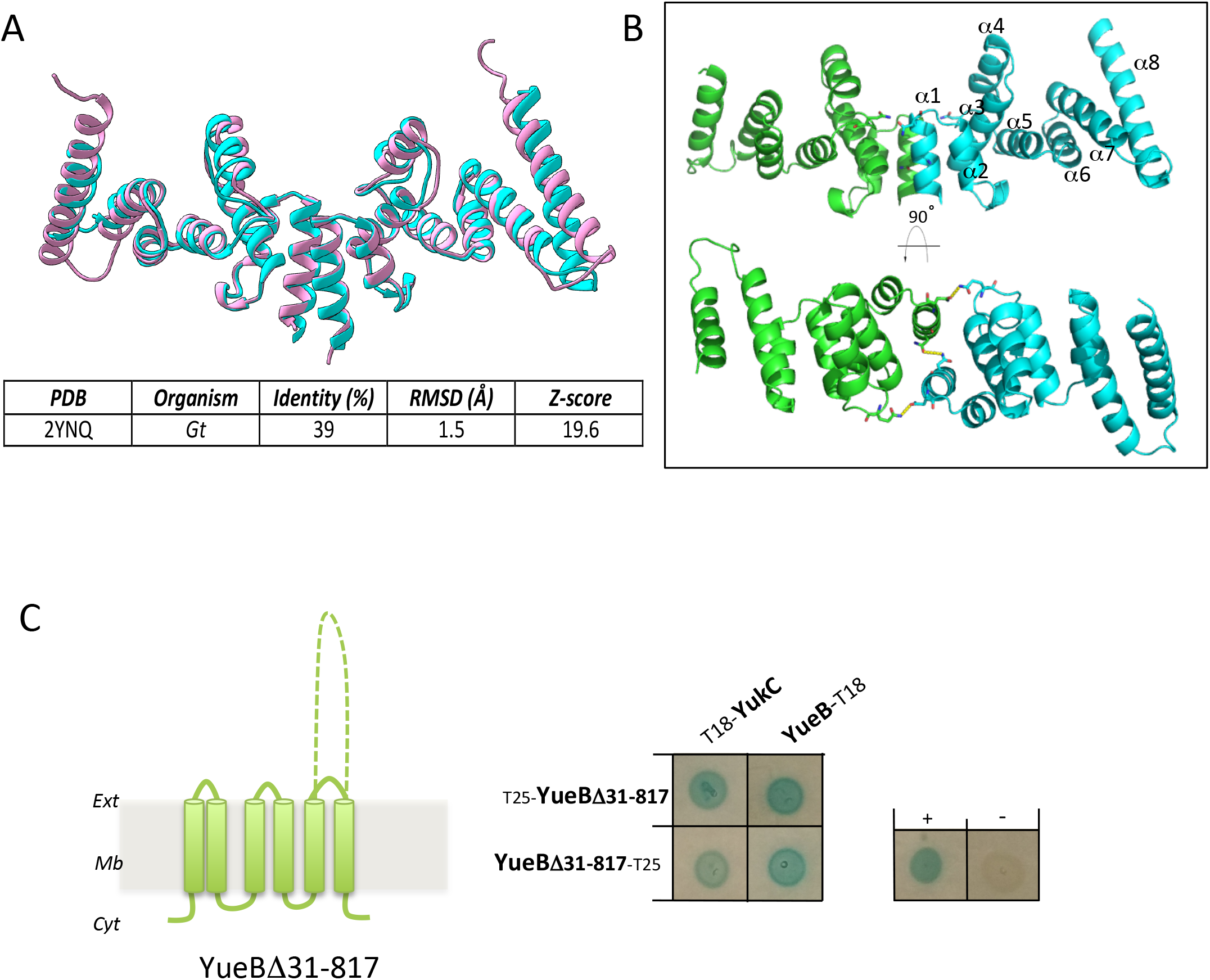
Analysis of the YukC_413_ extracellular region. **A)** Top: structural alignment of the extracellular domain of YukC_413_ (cyan) and EssB from *G. thermodenitrificans* (PDB 2YNQ, pink). Below: Table illustrating their structure similarities. **B)** Side and top views of the inter-molecular interactions between YukC_413_ monomers on the extracellular region. Numbering of the α-helices is indicated in the side view (top). **C)** Analysis of YukC-YueB interaction. Left: scheme of YueBΔ31-817 representing the large extracellular loop (dotted lines) mutated into a short loop (solid line). Right: BACTH assay on the interaction between YukC and YueBΔ31-817. The blue spots show interaction between YueBΔ31-817 and YukC Also, YueBΔ31-817 interaction with full-length YueB is shown. The positive and negative controls are displayed on the right.

**Figure 4-figure supplement 3.**
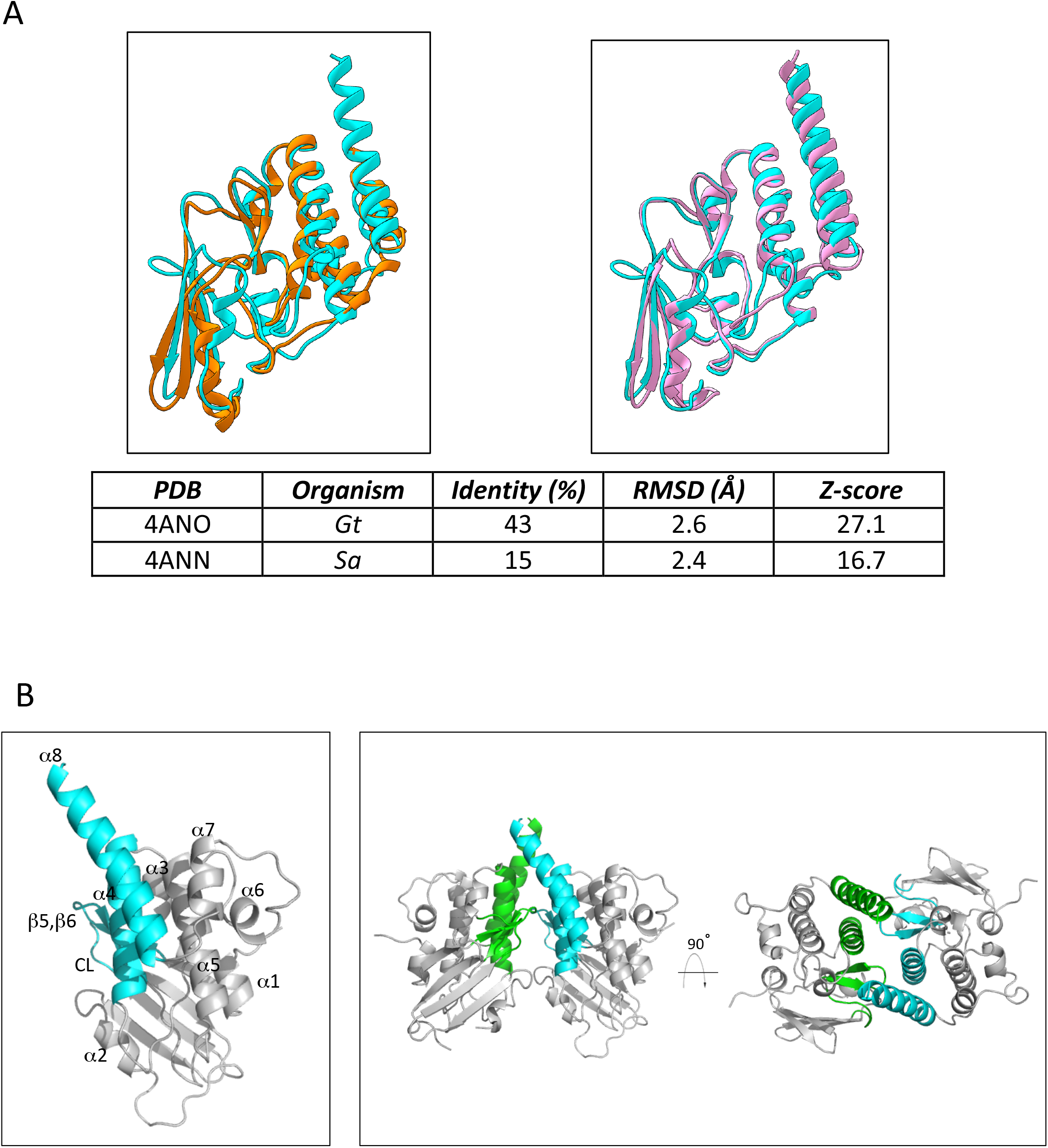
Comparison of the pseudokinase domains of YukC_413_ and *Sa*- and *Gt*- EssB.. **A)** Structural comparison between the pseudokinase domains of YukC_413_ (chains A, cyan) and the structures of the same region from *S. aureus* EssB (PDB 4ANN, orange and from *G. thermodenitrificans* (PDB 4ANO, pink). Below: Table illustrating their structure similarities. **B)** Interactions between the pseudokinase dimer. Only the regions involved in the intermolecular interactions are colored. Cyan: chain A. Green: chain B. Left panel: numbering of the α-helices in the pseudokinase domain seen (side view). The β-strands (β5 and β6) and the connection loop (CL) involved in the interaction are also indicated. Right panel: side and top views of the pseudokinase domains highlighting the interacting regions.

**Figure 4-figure supplement 4.**
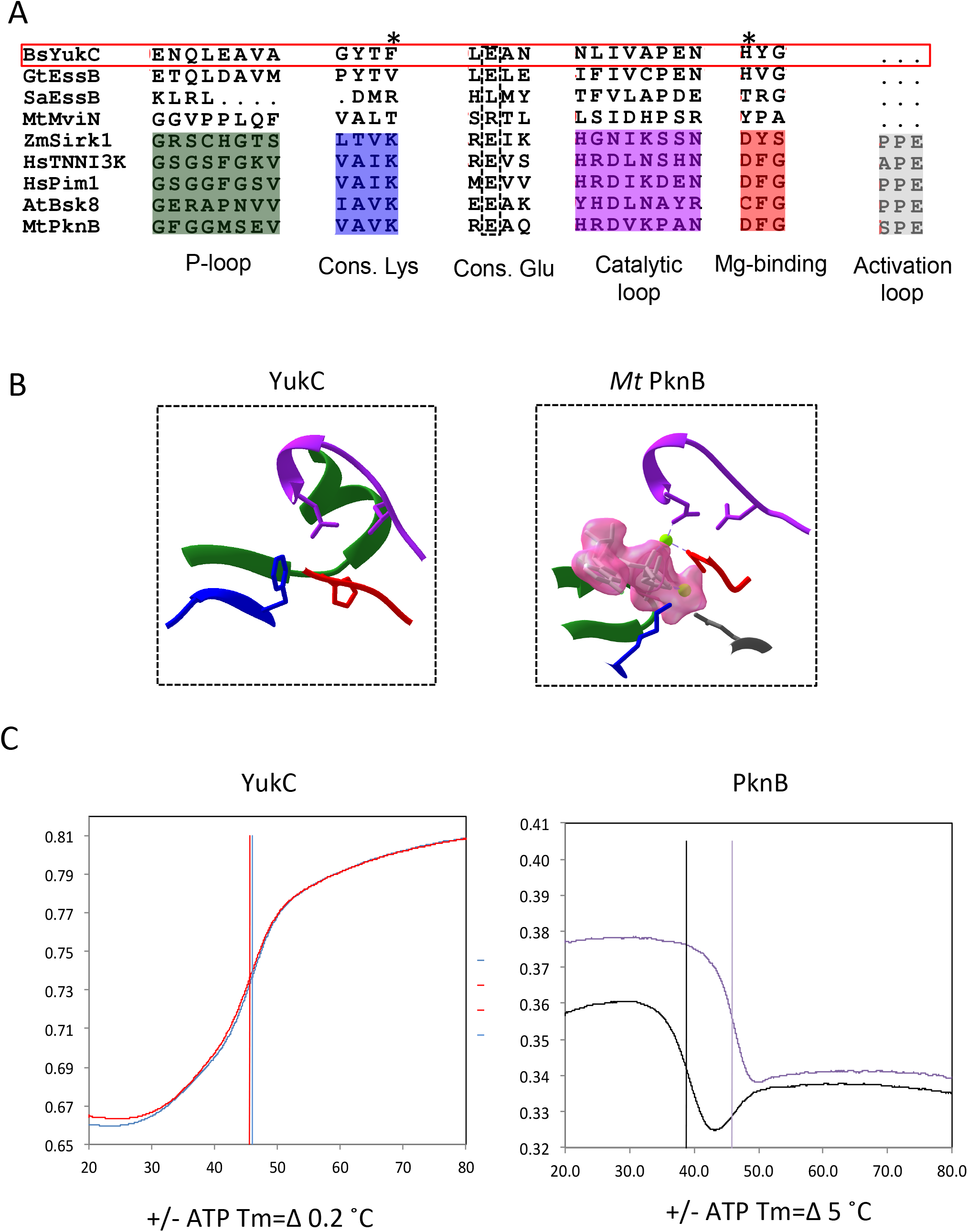
ATP is unable to bind to the YukC pseudokinase domain. **A)** Sequence alignment of *B. subtilis* YukC, its *S. aureus* and *G. thermodenitrificans* EssB homologs and known kinases and pseudokinases identified based on best matches using a DALI search. These include Sirk1, TNNI3K, Pim1, Bsk8 and *Mt*PknB (kinases) and the *Mt*MviN pseudokinase. The motifs characterizing the Hanks-type kinases are highlighted (such as the glycine-rich P-loop, the VAIK motif, the catalytic loop, the Mg^2+^-binding loop and the activation loop). **B)** Structural comparison between YukC_413_ and *Mt*PknB in complex with ATP (PDB: 1O6Y). The degenerated N-lobe YukC presents an additional α-helix in the ATP-binding pocket, with the large side-chain of F^26^ occluding the ATP binding pocket. **C)** Differential scanning fluorimetry (NanoDSF) measurement on YukC (left panel) and the kinase PknB from *M. tuberculosis* (right panel), in the presence and absence of ATP/Mg^2+^. The fluorescence signal, expressed as the 350 nm/330 nm emission ratio, is plotted as a function of the temperature. The average signal from three replicates is reported (two replicates were used in the case of PknB + ATP). The difference in the melting temperature (Tm) of each protein in the presence and in the absence of ATP, is indicated at the bottom of the respective plots.

**Figure 5-figure supplement 1.**
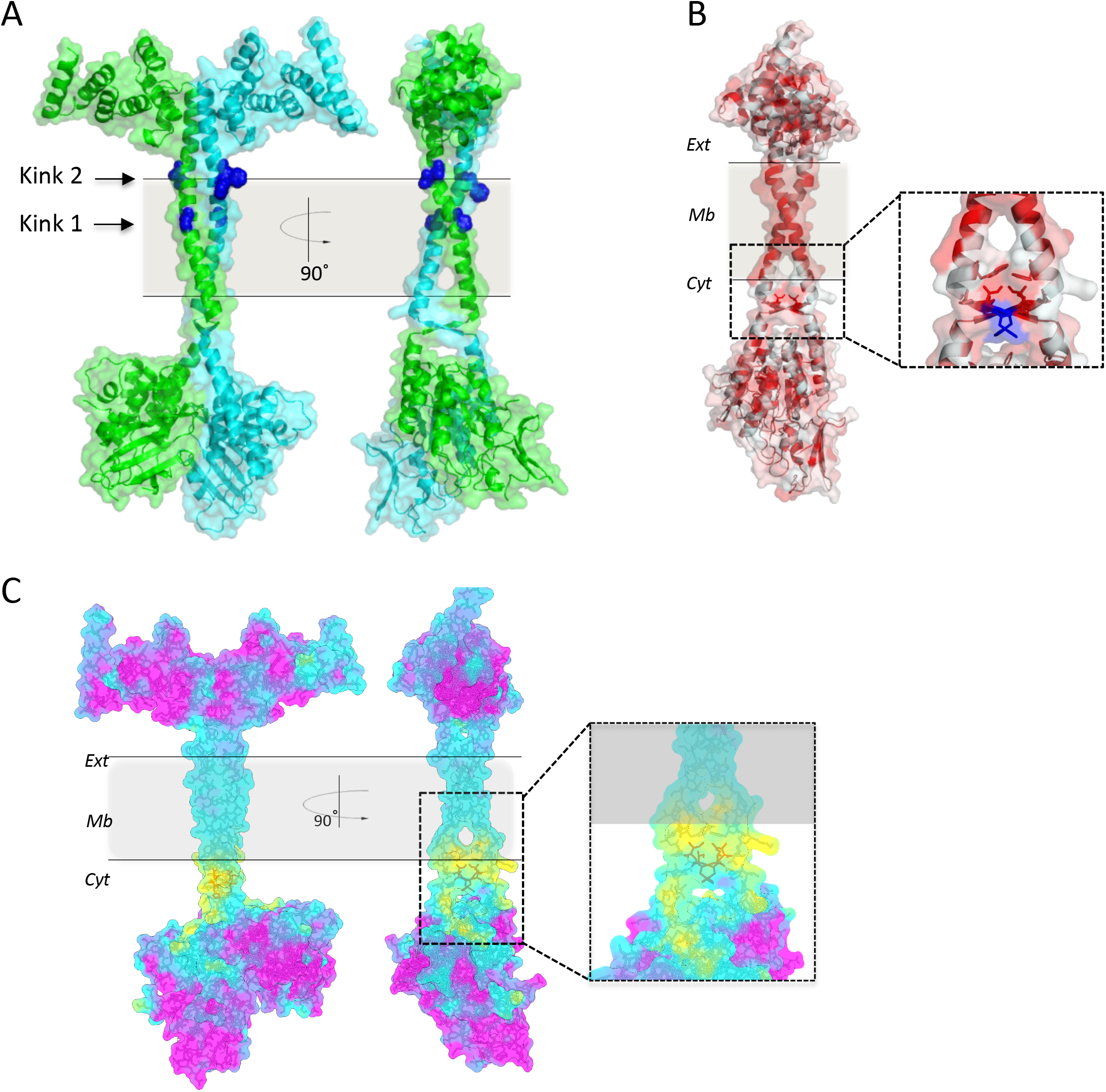
Analysis of the stalk and β-swap in the YukC_413_ dimer. **A)** YukC_413_ model representation showing kinks 1 and 2. Also the cavities inside the TM region and below are visible on the side view. The membrane region is always rendered in grey. **B)** Hydrophobicity residues in the YukC_413_ structure calculated by the PyMol software are colored in red. In the inset the hydrophobic patch over the β-swap is highlighted. The H^209^ is colored in blue. **C**) Electrostatic potential surface (violet: −10 kcal/(mole), cyan: 0 kcal/(mole), yellow: +10 kcal/(mole)) of the atomic model of YukC_413_, according to the coulombic law. In the inset the positive charges localized in proximity of the β-swap are highlighted and the residues I^208^, H^209^, I^210^ are colored in orange.

**Figure 5-figure supplement 2.**
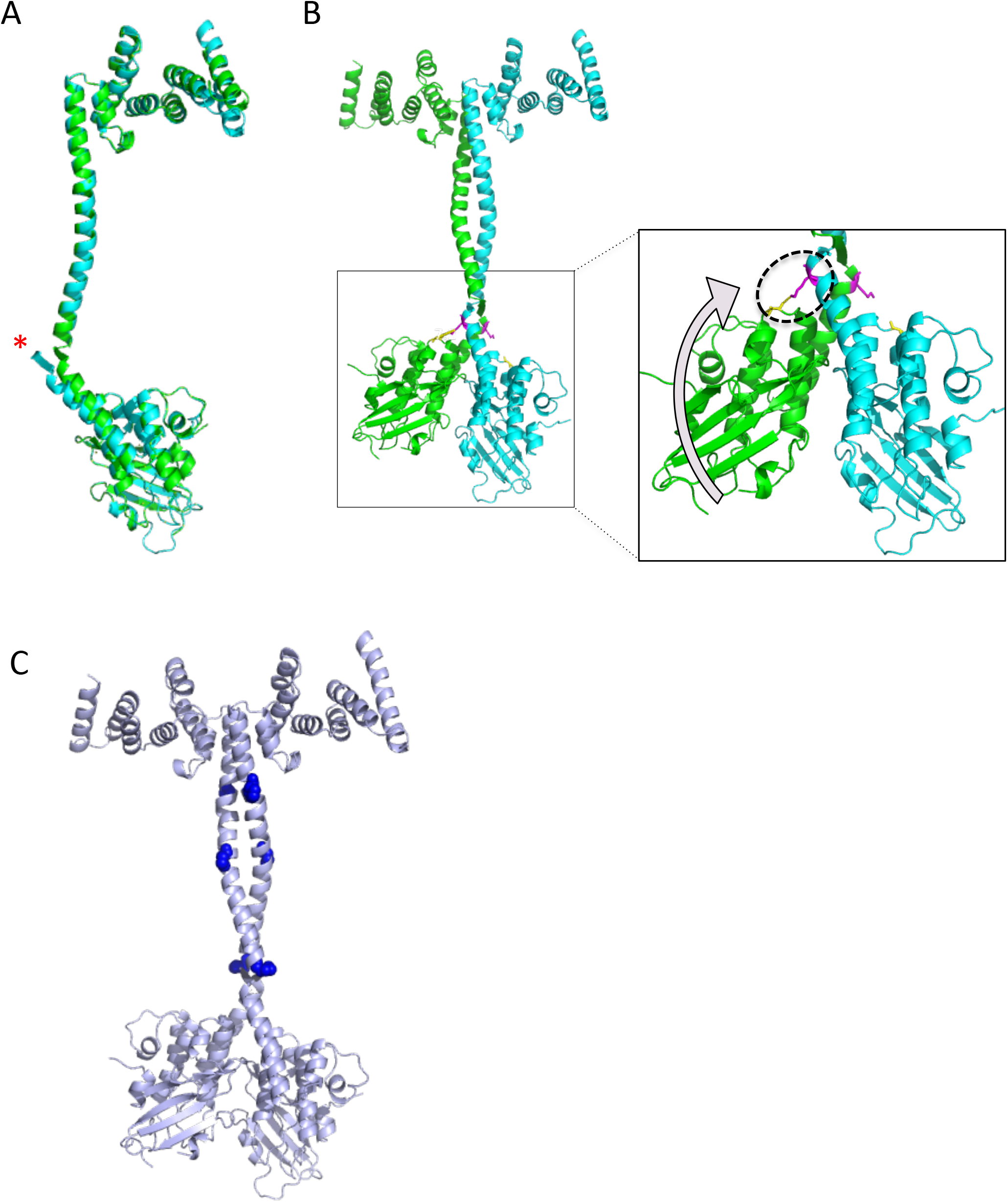
Structural basis of YukC asymmetry and prolines distribution. **A)** Alignment of chains A (cyan) and B (green). The orientation difference between the monomers is indicated by a red asterisk. **B)** Link between the pseudokinase domains different orientation and the asymmetrical interaction of K^205^ with A^80^. The K^205^ residues are depicted in magenta. In the inset: K^205^ - A^80^ interact in the monomer that is ‘lifted’ compared to the other. **C)** Proline residues P211, P231, P245 and P278 are depicted as blue spheres to highlight their localization along the YukC stalk.

**Table S1:**
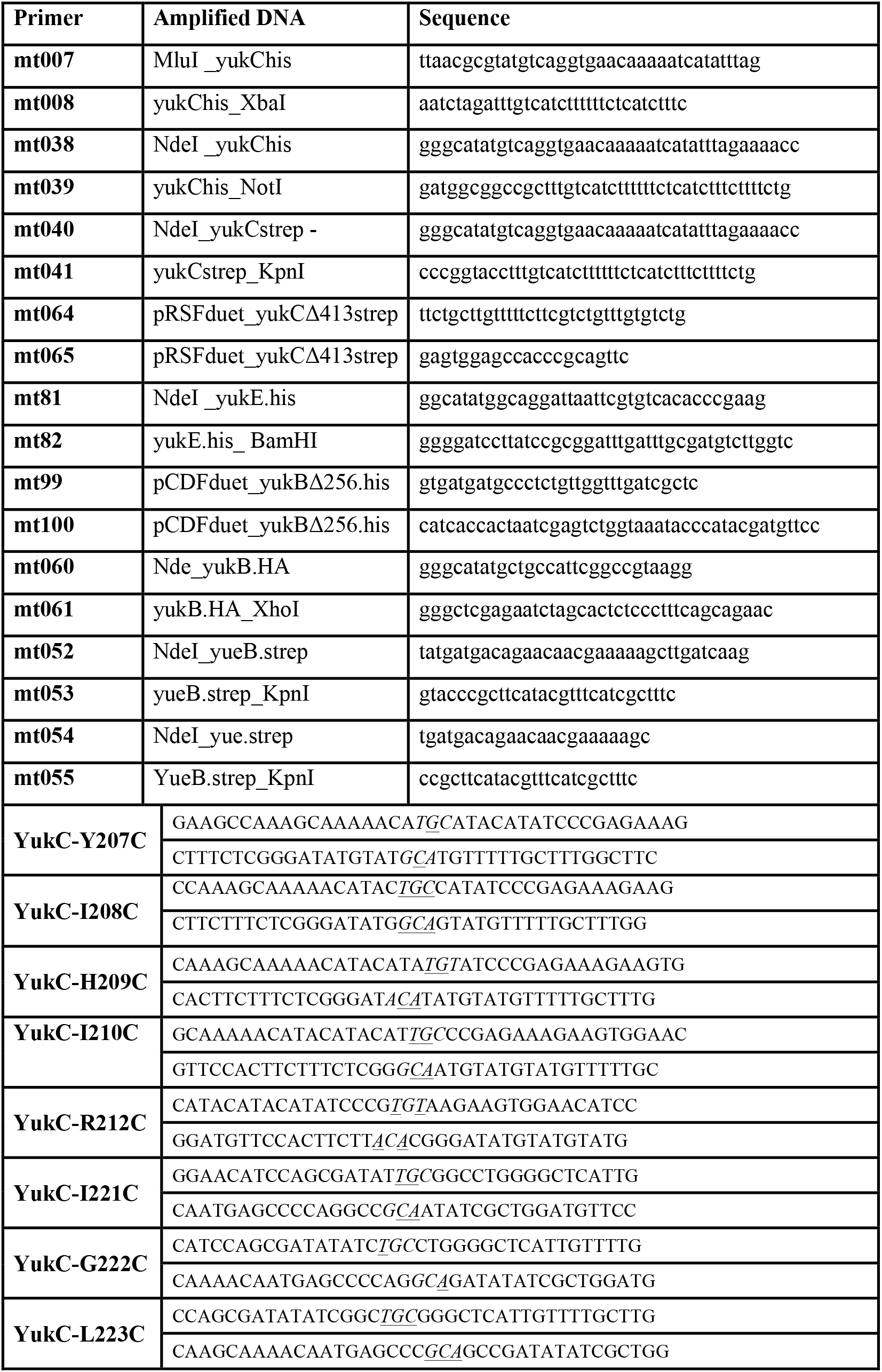
Primers used in this study

**Table S2:**
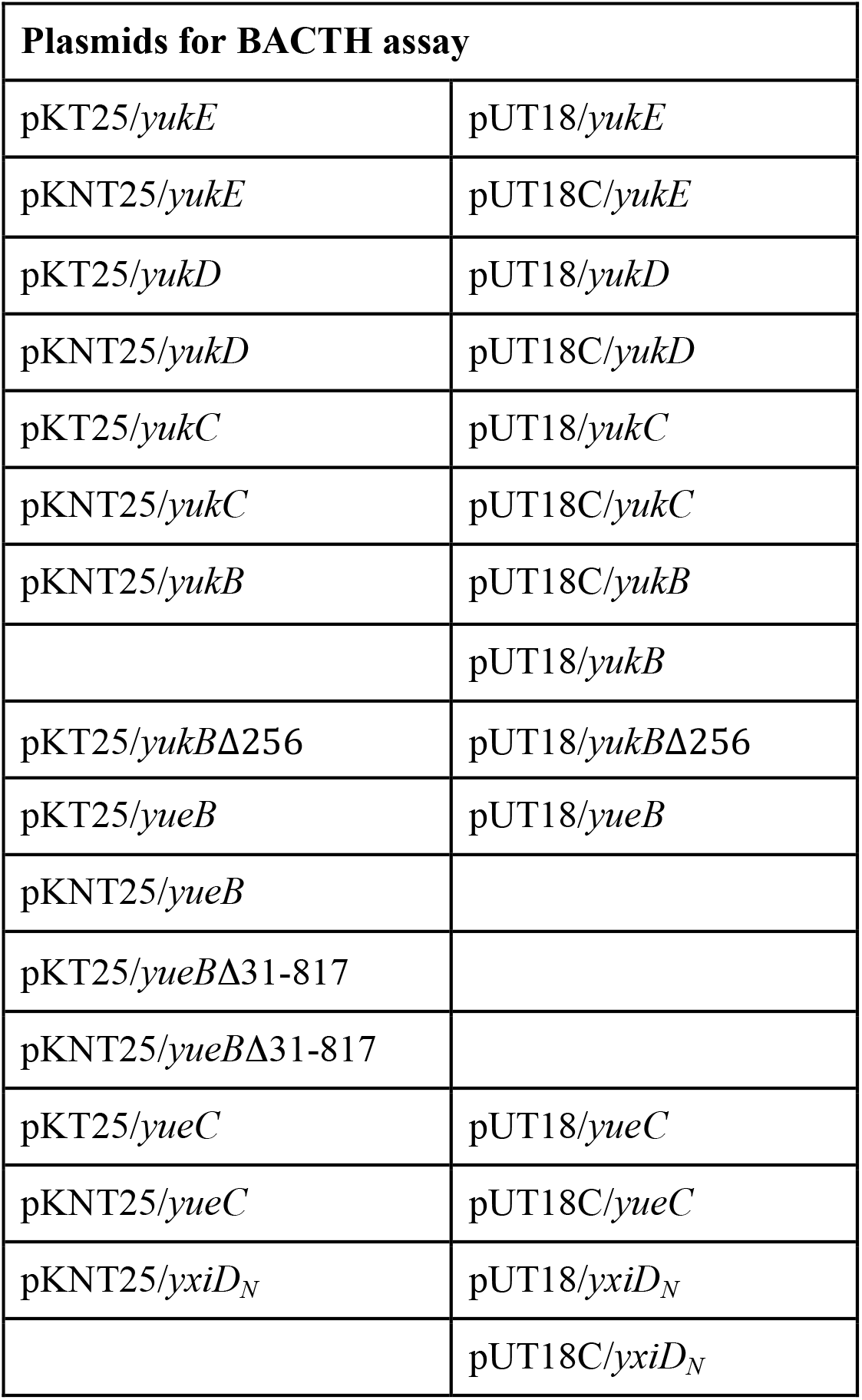
Plasmids used for BACTH assays

| Plasmids for BACTH assay |  |
| --- | --- |
| pKT25/*yukE* | pUT18/*yukE* |
| pKNT25/*yukE* | pUT18C/*yukE* |
| pKT25/*yukD* | pUT18/*yukD* |
| pKNT25/*yukD* | pUT18C/*yukD* |
| pKT25/*yukC* | pUT18/*yukC* |
| pKNT25/*yukC* | pUT18C/*yukC* |
| pKNT25/*yukB* | pUT18C/*yukB* |
|  | pUT18/*yukB* |
| pKT25/*yukB*Δ256 | pUT18/*yukB*Δ256 |
| pKT25/*yueB* | pUT18/*yueB* |
| pKNT25/*yueB* |  |
| pKT25/*yueB*Δ31-817 |  |
| pKNT25/*yueB*Δ31-817 |  |
| pKT25/*yueC* | pUT18/*yueC* |
| pKNT25/*yueC* | pUT18C/*yueC* |
| pKNT25/*yxiD<sub>N</sub>* | pUT18/*yxiD<sub>N</sub>* |
|  | pUT18C/*yxiD<sub>N</sub>* |

**Table S3:**
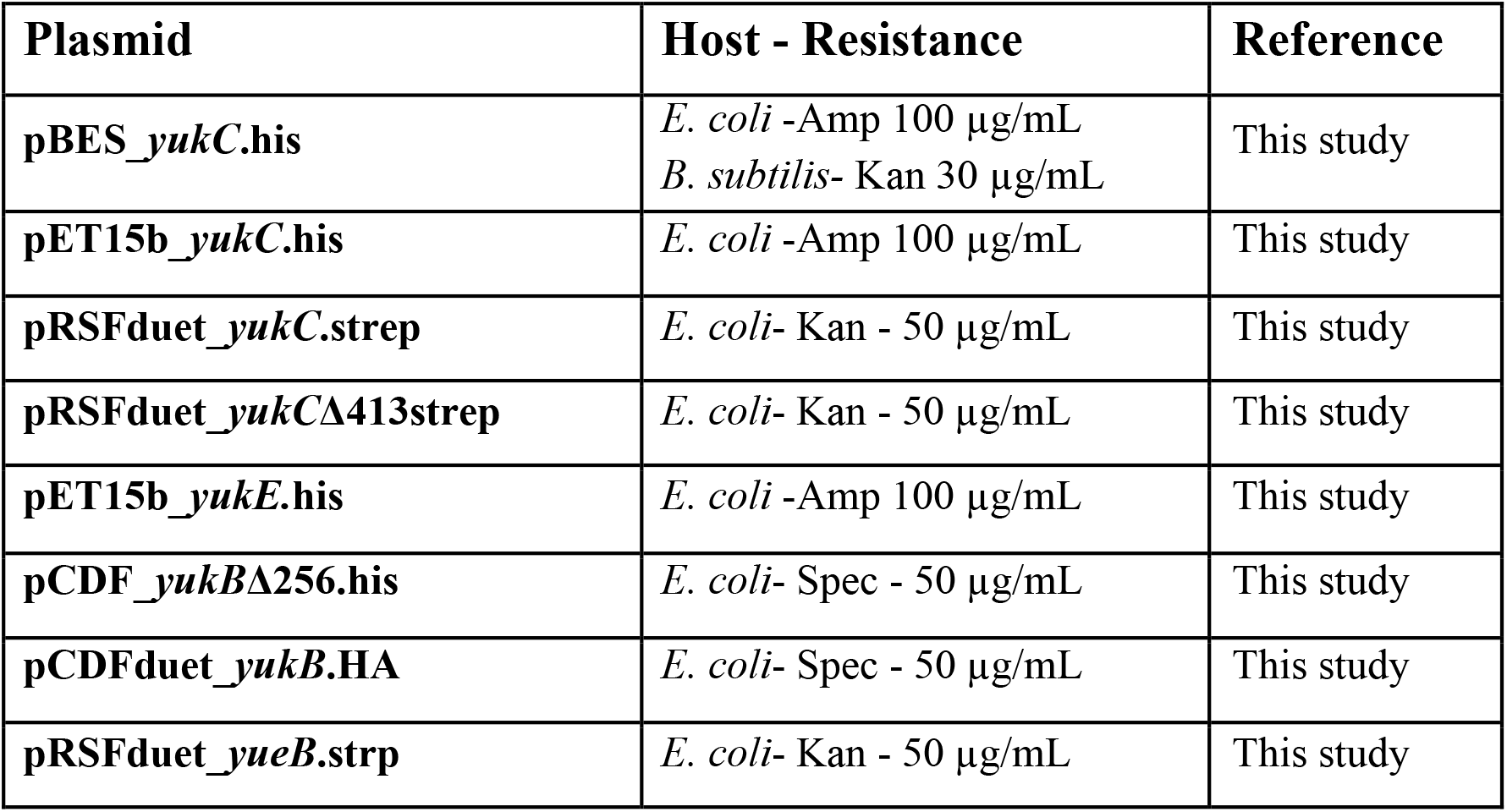
Plasmids used for protein expression

**Table S4:**
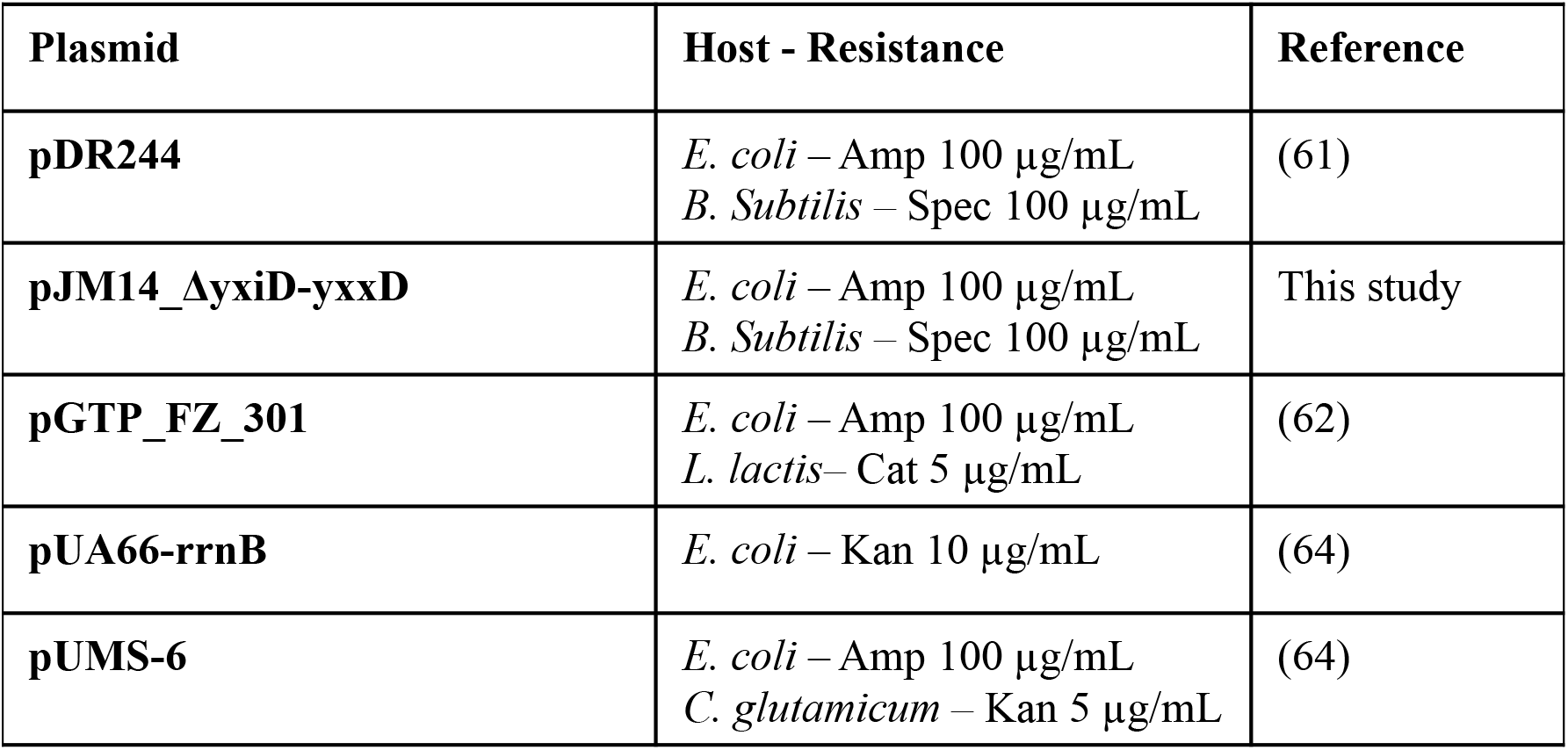
Plasmids used for competition assays

**Table S5:**
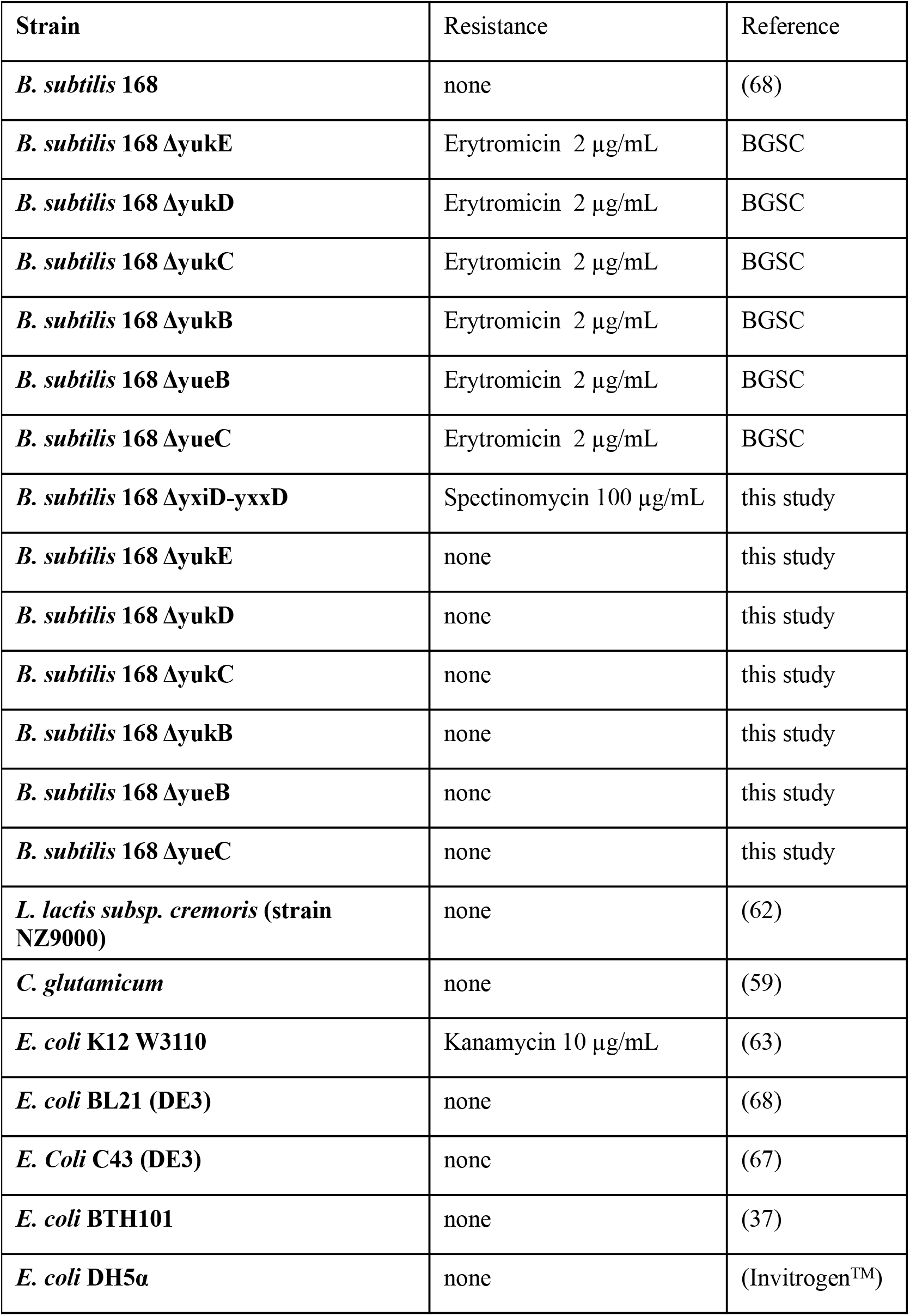
Strains used in this study

## Notes

### Competing Interest Statement

The authors have declared no competing interest.

## References

1. Groschel, M.I., Sayes, F., Simeone, R., Majlessi, L., and Brosch, R. (2016). ESX secretion systems: mycobacterial evolution to counter host immunity. Nature reviews. Microbiology.

2. Unnikrishnan, M., Constantinidou, C., Palmer, T., and Pallen, M.J. (2017). The Enigmatic Esx Proteins: Looking Beyond Mycobacteria. Trends Microbiol 25, 192–204.

3. Burts, M.L., Williams, W.A., DeBord, K., and Missiakas, D.M. (2005). EsxA and EsxB are secreted by an ESAT-6-like system that is required for the pathogenesis of Staphylococcus aureus infections. Proc Natl Acad Sci U S A 102, 1169–1174.

4. Garufi, G., Butler, E., and Missiakas, D. (2008). ESAT-6-Like Protein Secretion in Bacillus anthracis. Journal of bacteriology 190, 7004–7011.

5. Baptista, C., Barreto, H.C., and São-José, C. (2013). High levels of DegU-P activate an Esat-6-like secretion system in Bacillus subtilis. PloS one 8, e67840.

6. Rosenberg, O.S., Dovala, D., Li, X., Connolly, L., Bendebury, A., Finer-Moore, J., Holton, J., Cheng, Y., Stroud, R.M., and Cox, J.S. (2015). Substrates Control Multimerization and Activation of the Multi-Domain ATPase Motor of Type VII Secretion. Cell.

7. Poweleit, N., Czudnochowski, N., Nakagawa, R., Trinidad, D.D., Murphy, K.C., Sassetti, C.M., and Rosenberg, O.S. (2019). The structure of the endogenous ESX-3 secretion system. Elife 8.

8. Famelis, N., Rivera-Calzada, A., Degliesposti, G., Wingender, M., Mietrach, N., Skehel, J.M., Fernandez-Leiro, R., Bottcher, B., Schlosser, A., Llorca, O., et al. (2019). Architecture of the mycobacterial type VII secretion system. Nature.

9. Beckham, K.S., Ciccarelli, L., Bunduc, C.M., Mertens, H.D., Ummels, R., Lugmayr, W., Mayr, J., Rettel, M., Savitski, M.M., Svergun, D.I., et al. (2017). Structure of the mycobacterial ESX-5 type VII secretion system membrane complex by single-particle analysis. Nature microbiology 2, 17047.

10. Warne, B., Harkins, C.P., Harris, S.R., Vatsiou, A., Stanley-Wall, N., Parkhill, J., Peacock, S.J., Palmer, T., and Holden, M.T. (2016). The Ess/Type VII secretion system of Staphylococcus aureus shows unexpected genetic diversity. BMC genomics 17, 222.

11. Mietrach, N., Damian-Aparicio, D., Mielich-Suss, B., Lopez, D., and Geibel, S. (2020). Substrate Interaction with the EssC Coupling Protein of the Type VIIb Secretion System. J Bacteriol 202.

12. Zoltner, M., Ng, W.M., Money, J.J., Fyfe, P.K., Kneuper, H., Palmer, T., and Hunter, W.N. (2016). EssC: domain structures inform on the elusive translocation channel in the Type VII secretion system. Biochem J 473, 1941–1952.

13. Tanaka, Y., Kuroda, M., Yasutake, Y., Yao, M., Tsumoto, K., Watanabe, N., Ohta, T., and Tanaka, I. (2007). Crystal structure analysis reveals a novel forkhead-associated domain of ESAT-6 secretion system C protein in Staphylococcus aureus. Proteins 69, 659–664.

14. Jager, F., Kneuper, H., and Palmer, T. (2018). EssC is a specificity determinant for Staphylococcus aureus type VII secretion. Microbiology 164, 816–820.

15. Zoltner, M., Norman, D.G., Fyfe, P.K., El Mkami, H., Palmer, T., and Hunter, W.N. (2013). The architecture of EssB, an integral membrane component of the type VII secretion system. Structure 21, 595–603.

16. Zoltner, M., Fyfe, P.K., Palmer, T., and Hunter, W.N. (2013). Characterization of Staphylococcus aureus EssB, an integral membrane component of the Type VII secretion system: atomic resolution crystal structure of the cytoplasmic segment. Biochem J 449, 469–477.

17. Mahajan, A., Yuan, C., Lee, H., Chen, E.S., Wu, P.Y., and Tsai, M.D. (2008). Structure and function of the phosphothreonine-specific FHA domain. Science signaling 1, re12.

18. Ahmed, M.M., Aboshanab, K.M., Ragab, Y.M., Missiakas, D.M., and Aly, K.A. (2018). The transmembrane domain of the Staphylococcus aureus ESAT-6 component EssB mediates interaction with the integral membrane protein EsaA, facilitating partially regulated secretion in a heterologous host. Arch Microbiol 200, 1075–1086.

19. Aly, K.A., Anderson, M., Ohr, R.J., and Missiakas, D. (2017). Isolation of a membrane protein complex for type VII secretion in Staphylococcus aureus. J Bacteriol.

20. Bosserman, R.E., and Champion, P.A. (2017). Esx Systems and the Mycobacterial Cell Envelope: What’s the Connection? J Bacteriol 199.

21. Cao, Z., Casabona, M.G., Kneuper, H., Chalmers, J.D., and Palmer, T. (2016). The type VII secretion system of Staphylococcus aureus secretes a nuclease toxin that targets competitor bacteria. Nature microbiology 2, 16183.

22. Coros, A., Callahan, B., Battaglioli, E., and Derbyshire, K.M. (2008). The specialized secretory apparatus ESX-1 is essential for DNA transfer in Mycobacterium smegmatis. Mol Microbiol 69, 794–808.

23. Flint, J.L., Kowalski, J.C., Karnati, P.K., and Derbyshire, K.M. (2004). The RD1 virulence locus of Mycobacterium tuberculosis regulates DNA transfer in Mycobacterium smegmatis. Proc Natl Acad Sci U S A 101, 12598–12603.

24. Fyans, J.K., Bignell, D., Loria, R., Toth, I., and Palmer, T. (2013). The ESX/type VII secretion system modulates development, but not virulence, of the plant pathogen Streptomyces scabies. Mol Plant Pathol 14, 119–130.

25. Burts, M.L., DeDent, A.C., and Missiakas, D.M. (2008). EsaC substrate for the ESAT-6 secretion pathway and its role in persistent infections of Staphylococcus aureus. Mol Microbiol 69, 736–746.

26. Whitney, J.C., Peterson, S.B., Kim, J., Pazos, M., Verster, A.J., Radey, M.C., Kulasekara, H.D., Ching, M.Q., Bullen, N.P., Bryant, D., et al. (2017). A broadly distributed toxin family mediates contact-dependent antagonism between grampositive bacteria. Elife 6.

27. Zhang, D., Iyer, L.M., and Aravind, L. (2011). A novel immunity system for bacterial nucleic acid degrading toxins and its recruitment in various eukaryotic and DNA viral systems. Nucleic Acids Res 39, 4532–4552.

28. Holberger, L.E., Garza-Sanchez, F., Lamoureux, J., Low, D.A., and Hayes, C.S. (2012). A novel family of toxin/antitoxin proteins in Bacillus species. FEBS letters 586, 132–136.

29. Hayes, C.S., Aoki, S.K., and Low, D.A. (2010). Bacterial contact-dependent delivery systems. Annu Rev Genet 44, 71–90.

30. Huppert, L.A., Ramsdell, T.L., Chase, M.R., Sarracino, D.A., Fortune, S.M., and Burton, B.M. (2014). The ESX system in Bacillus subtilis mediates protein secretion. PLoS One 9, e96267.

31. Sao-Jose, C., Baptista, C., and Santos, M.A. (2004). Bacillus subtilis operon encoding a membrane receptor for bacteriophage SPP1. J Bacteriol 186, 8337–8346.

32. Bachmann, B.J. (1972). Pedigrees of some mutant strains of Escherichia coli K-12. Bacteriol Rev 36, 525–557.

33. Kato, T., Maeda, K., Kasuya, H., and Matsuda, T. (1999). Complete Growth Inhibition of Bacillus subtilis by Nisin-Producing Lactococci in Fermented Soybeans. Biosci Biotechnol Biochem 63, 642–647.

34. Alegria, A., Delgado, S., Roces, C., Lopez, B., and Mayo, B. (2010). Bacteriocins produced by wild Lactococcus lactis strains isolated from traditional, starter-free cheeses made of raw milk. Int J Food Microbiol 143, 61–66.

35. Jager, F., Zoltner, M., Kneuper, H., Hunter, W.N., and Palmer, T. (2016). Membrane interactions and self-association of components of the Ess/Type VII secretion system of Staphylococcus aureus. FEBS letters 590, 349–357.

36. Aly, K.A., Anderson, M., Ohr, R.J., and Missiakas, D. (2017). Isolation of a Membrane Protein Complex for Type VII Secretion in Staphylococcus aureus. J Bacteriol 199.

37. Karimova, G., Pidoux, J., Ullmann, A., and Ladant, D. (1998). A bacterial two-hybrid system based on a reconstituted signal transduction pathway. Proc Natl Acad Sci U S A 95, 5752–5756.

38. Sysoeva, T.A., Zepeda-Rivera, M.A., Huppert, L.A., and Burton, B.M. (2014). Dimer recognition and secretion by the ESX secretion system in Bacillus subtilis. Proc Natl Acad Sci U S A 111, 7653–7658.

39. Holm, L., and Rosenstrom, P. (2010). Dali server: conservation mapping in 3D. Nucleic Acids Res 38, W545–549.

40. Ortiz-Lombardia, M., Pompeo, F., Boitel, B., and Alzari, P.M. (2003). Crystal structure of the catalytic domain of the PknB serine/threonine kinase from Mycobacterium tuberculosis. The Journal of biological chemistry 278, 13094–13100.

41. Griffin, J.E., Gawronski, J.D., Dejesus, M.A., Ioerger, T.R., Akerley, B.J., and Sassetti, C.M. (2011). High-resolution phenotypic profiling defines genes essential for mycobacterial growth and cholesterol catabolism. PLoS Pathog 7, e1002251.

42. Krissinel, E., and Henrick, K. (2007). Inference of macromolecular assemblies from crystalline state. Journal of molecular biology 372, 774–797.

43. Patel, P.S., Huang, S., Fisher, S., Pirnik, D., Aklonis, C., Dean, L., Meyers, E., Fernandes, P., and Mayerl, F. (1995). Bacillaene, a novel inhibitor of procaryotic protein synthesis produced by Bacillus subtilis: production, taxonomy, isolation, physico-chemical characterization and biological activity. J Antibiot (Tokyo) 48, 997–1003.

44. Wright, G.D. (2005). Bacterial resistance to antibiotics: enzymatic degradation and modification. Adv Drug Deliv Rev 57, 1451–1470.

45. Lopez, D., Vlamakis, H., Losick, R., and Kolter, R. (2009). Paracrine signaling in a bacterium. Genes & development 23, 1631–1638.

46. Hoefler, B.C., Gorzelnik, K.V., Yang, J.Y., Hendricks, N., Dorrestein, P.C., and Straight, P.D. (2012). Enzymatic resistance to the lipopeptide surfactin as identified through imaging mass spectrometry of bacterial competition. Proc Natl Acad Sci U S A 109, 13082–13087.

47. Straight, P.D., Willey, J.M., and Kolter, R. (2006). Interactions between Streptomyces coelicolor and Bacillus subtilis: Role of surfactants in raising aerial structures. J Bacteriol 188, 4918–4925.

48. Koskiniemi, S., Lamoureux, J.G., Nikolakakis, K.C., t’Kint de Roodenbeke, C., Kaplan, M.D., Low, D.A., and Hayes, C.S. (2013). Rhs proteins from diverse bacteria mediate intercellular competition. Proc Natl Acad Sci U S A 110, 7032–7037.

49. Jamet, A., Charbit, A., and Nassif, X. (2018). Antibacterial Toxins: Gram-Positive Bacteria Strike Back! Trends Microbiol 26, 89–91.

50. Baptista, C., Barreto, H.C., and Sao-Jose, C. (2013). High levels of DegU-P activate an Esat-6-like secretion system in Bacillus subtilis. PLoS One 8, e67840.

51. Durocher, D., and Jackson, S.P. (2002). The FHA domain. FEBS letters 513, 58–66.

52. Pompeo, F., Byrne, D., Mengin-Lecreulx, D., and Galinier, A. (2018). Dual regulation of activity and intracellular localization of the PASTA kinase PrkC during Bacillus subtilis growth. Scientific reports 8, 1660.

53. Stancik, I.A., Sestak, M.S., Ji, B., Axelson-Fisk, M., Franjevic, D., Jers, C., Domazet-Loso, T., and Mijakovic, I. (2018). Serine/Threonine Protein Kinases from Bacteria, Archaea and Eukarya Share a Common Evolutionary Origin Deeply Rooted in the Tree of Life. Journal of molecular biology 430, 27–32.

54. Lee, Y., Nishizawa, T., Yamashita, K., Ishitani, R., and Nureki, O. (2015). Structural basis for the facilitative diffusion mechanism by SemiSWEET transporter. Nature communications 6, 6112.

55. Jacob, J., Duclohier, H., and Cafiso, D.S. (1999). The role of proline and glycine in determining the backbone flexibility of a channel-forming peptide. Biophysical journal 76, 1367–1376.

56. Chow, W.Y., Forman, C.J., Bihan, D., Puszkarska, A.M., Rajan, R., Reid, D.G., Slatter, D.A., Colwell, L.J., Wales, D.J., Farndale, R.W., et al. (2018). Proline provides site-specific flexibility for in vivo collagen. Scientific reports 8, 13809.

57. Kumeta, M., Konishi, H.A., Zhang, W., Sakagami, S., and Yoshimura, S.H. (2018). Prolines in the alpha-helix confer the structural flexibility and functional integrity of importin-beta. J Cell Sci 131.

58. Fernandez, P., Porrini, L., Albanesi, D., Abriata, L.A., Dal Peraro, M., de Mendoza, D., and Mansilla, M.C. (2019). Transmembrane Prolines Mediate Signal Sensing and Decoding in Bacillus subtilis DesK Histidine Kinase. MBio 10.

59. Mielich-Suss, B., Wagner, R.M., Mietrach, N., Hertlein, T., Marincola, G., Ohlsen, K., Geibel, S., and Lopez, D. (2017). Flotillin scaffold activity contributes to type VII secretion system assembly in Staphylococcus aureus. PLoS Pathog 13, e1006728.

60. Lemon, K.P., and Grossman, A.D. (1998). Localization of bacterial DNA polymerase: evidence for a factory model of replication. Science 282, 1516–1519.

61. Koo, B.M., Kritikos, G., Farelli, J.D., Todor, H., Tong, K., Kimsey, H., Wapinski, I., Galardini, M., Cabal, A., Peters, J.M., et al. (2017). Construction and Analysis of Two Genome-Scale Deletion Libraries for Bacillus subtilis. Cell Syst 4, 291–305 e297.

62. Tremillon, N., Issaly, N., Mozo, J., Duvignau, T., Ginisty, H., Devic, E., and Poquet, I. (2010). Production and purification of staphylococcal nuclease in Lactococcus lactis using a new expression-secretion system and a pH-regulated mini-reactor. Microb Cell Fact 9, 37.

63. Santin, Y.G., Doan, T., Lebrun, R., Espinosa, L., Journet, L., and Cascales, E. (2018). In vivo TssA proximity labelling during type VI secretion biogenesis reveals TagA as a protein that stops and holds the sheath. Nature microbiology 3, 1304–1313.

64. Sogues, A., Martinez, M., Gaday, Q., Ben Assaya, M., Grana, M., Voegele, A., VanNieuwenhze, M., England, P., Haouz, A., Chenal, A., et al. (2020). Essential dynamic interdependence of FtsZ and SepF for Z-ring and septum formation in Corynebacterium glutamicum. Nature communications 11, 1641.

65. Bindels, D.S., Haarbosch, L., van Weeren, L., Postma, M., Wiese, K.E., Mastop, M., Aumonier, S., Gotthard, G., Royant, A., Hink, M.A., et al. (2017). mScarlet: a bright monomeric red fluorescent protein for cellular imaging. Nat Methods 14, 53–56.

66. Schindelin, J., Arganda-Carreras, I., Frise, E., Kaynig, V., Longair, M., Pietzsch, T., Preibisch, S., Rueden, C., Saalfeld, S., Schmid, B., et al. (2012). Fiji: an open-source platform for biological-image analysis. Nat Methods 9, 676–682.

67. Miroux, B., and Walker, J.E. (1996). Over-production of proteins in Escherichia coli: mutant hosts that allow synthesis of some membrane proteins and globular proteins at high levels. Journal of molecular biology 260, 289–298.

68. Wood, W.B. (1966). Host specificity of DNA produced by Escherichia coli: bacterial mutations affecting the restriction and modification of DNA. Journal of molecular biology 16, 118–133.

69. Vonrhein, C., Flensburg, C., Keller, P., Sharff, A., Smart, O., Paciorek, W., Womack, T., and Bricogne, G. (2011). Data processing and analysis with the autoPROC toolbox. Acta Crystallogr D Biol Crystallogr 67, 293–302.

70. McCoy, A.J., Grosse-Kunstleve, R.W., Adams, P.D., Winn, M.D., Storoni, L.C., and Read, R.J. (2007). Phaser crystallographic software. J Appl Crystallogr 40, 658–674.

71. Williams, C.J., Headd, J.J., Moriarty, N.W., Prisant, M.G., Videau, L.L., Deis, L.N., Verma, V., Keedy, D.A., Hintze, B.J., Chen, V.B., et al. (2018). MolProbity: More and better reference data for improved all-atom structure validation. Protein science: a publication of the Protein Society 27, 293–315.

72. Terwilliger, T.C., Grosse-Kunstleve, R.W., Afonine, P.V., Moriarty, N.W., Zwart, P.H., Hung, L.W., Read, R.J., and Adams, P.D. (2008). Iterative model building, structure refinement and density modification with the PHENIX AutoBuild wizard. Acta Crystallogr D Biol Crystallogr 64, 61–69.

73. Emsley, P., Lohkamp, B., Scott, W.G., and Cowtan, K. (2010). Features and development of Coot. Acta Crystallogr D Biol Crystallogr 66, 486–501.

74. Smart, O.S., Womack, T.O., Flensburg, C., Keller, P., Paciorek, W., Sharff, A., Vonrhein, C., and Bricogne, G. (2012). Exploiting structure similarity in refinement: automated NCS and target-structure restraints in BUSTER. Acta Crystallogr D Biol Crystallogr 68, 368–380.

75. Notredame, C., Higgins, D.G., and Heringa, J. (2000). T-Coffee: A novel method for fast and accurate multiple sequence alignment. Journal of molecular biology 302, 205–217.

76. Pettersen, E.F., Goddard, T.D., Huang, C.C., Couch, G.S., Greenblatt, D.M., Meng, E.C., and Ferrin, T.E. (2004). UCSF Chimera--a visualization system for exploratory research and analysis. J Comput Chem 25, 1605–1612.

